# Brain mechanism of foraging: reward-dependent synaptic plasticity or neural integration of values?

**DOI:** 10.1101/2022.09.25.509030

**Authors:** Ulises Pereira-Obilinovic, Han Hou, Karel Svoboda, Xiao-Jing Wang

**Affiliations:** Center for Neural Science, New York University, New York, NY 10003, USA; Allen Institute for Neural Dynamics, Seattle, WA 98109, USA

**Keywords:** Foraging, Value-based decision-making, Reinforcement learning, Neural integrator

## Abstract

During foraging behavior, action values are persistently encoded in neural activity and updated depending on the history of choice outcomes. What is the neural mechanism for action value maintenance and updating? Here we explore two contrasting network models: synaptic learning of action value versus neural integration. We show that both models can reproduce extant experimental data, but they yield distinct predictions about the underlying biological neural circuits. In particular, the neural integrator model but not the synaptic model requires that reward signals are mediated by neural pools selective for action alternatives and their projections are aligned with linear attractor axes in the valuation system. We demonstrate experimentally observable neural dynamical signatures and feasible perturbations to differentiate the two contrasting scenarios, suggesting that the synaptic model is a more robust candidate mechanism. Overall, this work provides a modeling framework to guide future experimental research on probabilistic foraging.

---

**D**uring foraging, action values are stored in short-term memory over times of several behavioral trials, much longer than intrinsic neural time scales. The stored action values bias actions to increase reward acquisition. Action values are updated depending on the history of choices and rewards. Single-neuron activity in the monkey posterior parietal cortex (1, 2), the mouse medial prefrontal (mPFC) (3) and retrosplenial cortex (RSC) (4), is correlated with estimated action values. The activity of some of these neurons is persistent, lasting for seconds during the entire inter-trial interval (ITI) and updated after the trial ends (3, 4).

Such value encoding and update during foraging has traditionally been explained by reward-dependent synaptic plasticity (5, 6). In these models, Hebbian plasticity combined with reward signals (7) modifies synapses that connect feedforward inputs to a recurrent network that displays winner-take-all dynamics. Action values are encoded in the feedforward synaptic weights to action-encoding selective populations, biasing the network’s winner-take-all dynamics. However, this stands in contrast with recordings of neurons in the mice mPFC and RSC that display graded persistent encoding of action value (3, 4).

Artificial neural networks (ANNs) can be trained to produce foraging behavior without synaptic changes in synaptic weights (8). These ANNs incorporate mechanisms referred to as *g*ates with unknown neuronal bases which enables them to maintain signals in memory for extended periods (9, 10). It remains uncertain whether foraging without changes in synaptic weights represents a robust principle underlying foraging in the brain. However, these findings in conjunction with the observed persistent graded activity proportional to action values (3, 4) have raised an interesting alternative mechanism for foraging without synaptic changes by virtue of neural integration such as what is found in line attractor models (11, 12).

In this work, we investigated these two contrasting mechanisms for foraging: reward-dependent synaptic plasticity and neural integration of values which does not rely on synaptic weight changes. We build biologically plausible yet simple models based on the two above scenarios. Both recapitulate the behavior and important features of neural dynamics during foraging (3, 4). However, they differ qualitatively in their network architecture, dynamical properties, and response to perturbations. We found the synaptic mechanism is much more robust to perturbations in the neural activity and the connectivity than the neural integration of values. The neural integrator model requires vectorial reward prediction error (RPE) signals in an *on-manifold* alignment with the line attractor axes. Lastly, we explored structured foraging tasks which it is advantageous anti-correlated action value updates. We predict that in these tasks the neural integrator requires selective changes in the vectorial RPE alignment, whereas the synaptic model accommodates these updates by unselective changes of shared inhibition. We outline experimental predictions that can be used to disprove these theories using current neurotechnologies. Our work provides a modeling framework to investigate the neural mechanisms underlying foraging behavior.

## Results

### Two mechanisms for value maintenance and update

In the neural integrator model (Fig. 1A), action values are stored in directions in the neural space of slow variation in population activity or line attractor axes. The neural integrator integrates rewards by updating action values by an input proportional to the reward prediction error (RPE) in the direction of action value encoding (see Fig. 1A). In this mechanism, changes in the synaptic efficacy are not needed for action value update, similar to a theoretical proposal in RNN-based agents (8). On the other hand, in the synaptic model (Fig. 1C), a scalar feedback triggered by reward enables Hebbian plasticity on the recurrent connections of the valuation network, in turn modifying in a reward-dependent manner the synaptic efficacy on the network’s recurrent connections (5, 6). Since synaptic inputs drive spiking activity, action values modulate neuronal activity. Action values are maintained in neural space in a single fixed-point attractor that shifts its location on a trial-by-trial basis (Fig. 1C).

**Fig. 1.**
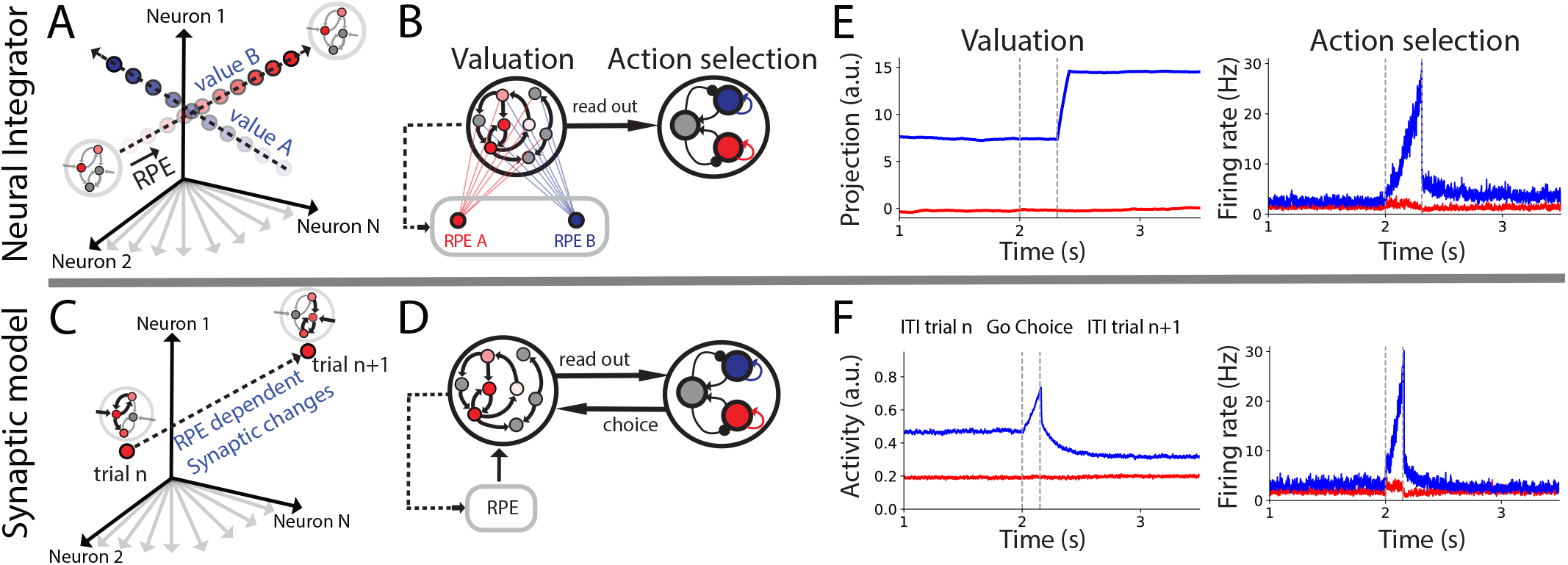
Neural integrator (Top row) vs. synaptic model (bottom row) for foraging behavior. (A) Maintainance and neural integration of values. The value for the two contingencies is maintained in two different line attractors embedded in high-dimensional neural space. Increasing color opacity represents increasing value. After the reward is collected in the task, RPE signals align to the line attractor corresponding to the chosen contingency update values by stirring the neural activity along the corresponding line attractor. Changes in value are not caused by changes in synaptic weights. (B) Network architecture for the neural integrator. Action value is maintained in the valuation network. These values are read out by the action selection network for producing the choice through a winner-take-all dynamics. RPE signals are computed by subtracting from the trial-by-trial reward the action value encoded in the valuation network (see dashed line arrow). The RPE signal is routed in a contingency-dependence fashion to the valuation network to update action value. (C) Maintenance and integration of values through synaptic plasticity. The value of the two contingencies is maintained in neural space in a single fixed-point attractor. Changes in synaptic weights shift the location of the fixed-point attractor in neural space on a trial-by-trial basis. (D) Network architecture for the synaptic model. Values are maintained by two selective populations in the valuation network. As in (B), values are read out by the action selection network for producing the choice. Feedback choice signals elevate the activity of selective populations in the valuation network. RPE signals are computed by subtracting from the trial-by-trial reward the action value encoded in the valuation network (see dashed line arrow). The RPE is combined with elevated activity to produce synaptic plasticity. The dynamics of one trial for the valuation (left) and action selection (right) networks for the Neural integrator (E) and synaptic model (F).

We explore these two contrasting neural mechanisms underlying action value maintenance and its reward-dependent update during foraging. We develop circuit models for the two mechanisms that allow us to explore experimental tests for these two mechanisms. These models were guided by anatomical and physiological evidence of the neural circuits involved in foraging behavior (3, 13). We aim for these models to be simple enough to be suitable for mechanistic understanding and mathematical analysis, but to include key details to the extent their predictions can be interpreted in actual brain circuits.

We first focus on the dynamic foraging task (DFT) (2–4, 14, 15) (see Methods). Briefly, in this task, after a go cue, the subject freely chooses among two contingencies that deliver reward with nonstationary probabilities. The reward probability for the two contingencies changes in blocks of tens to hundreds of trials. On each trial, a go cue instructs the subject to make a choice and collect reward. After a reward is delivered probabilistically, the subject waits an ITI of several seconds after the next go cue. Different from the two-armed bandit task (16), the reward is baited (2–4, 14, 15) in this task: once the reward is assigned, it remains assigned to the contingency until its consumption even if the contingency is not chosen.

The network architecture consists of two inter-connected areas, each modeled as recurrent networks (Fig. 1B and D). Action values for the two contingencies are maintained and updated depending on reward in the ‘valuation network’. Actions are selected depending on action values in the ‘action selection network’. This architecture is inspired by the separation between the limbic and motor information streams during decision-making (13, 17).

In the neural integrator model, action values for the two contingencies are maintained by the population activity in two linear axes embedded in a high-dimensional neural space (the space spanned by the activity of all neurons in the network) (Fig. 1A) that we refer as line attractor axes. After the action is selected and the reward is delivered, an RPE signal updates the corresponding action value represented in the line attractor axes (Fig. 1B). Importantly, although the two line attractor axes are orthogonal in neural space, the network connectivity is recurrent, and overlapping populations encode action values for the two contingencies (Fig. S1 in SI). Two predictions can be derived directly from our neural integrator model’s assumptions:

1. RPE signals have to be at least partially aligned to the action value encoding axis corresponding to the chosen contingency to produce sizable changes in value (see more in the section response to perturbations). Therefore, unlike the view in which the RPE is a global scalar signal (18), for the neural integrator model, RPE must project to a selective neural population in a choice-dependent manner to update a particular action value in any given trial.
2. For choice signals from the action selection network to not disrupt the value encoding in the valuation networks we expect feedback to be weak or approximately orthogonal to the line attractor axes.

In the synaptic model, the recurrent synaptic weights in the valuation network are updated according to the three-factor learning rule (see Eq. (17)). After the action is selected, the pre and post-synaptic activity of chosen selective populations in the valuation network is transiently elevated after the action is selected due to the feedback from the action selection network (Fig. 1B and D). In contrast, little to no feedback activity is produced in the corresponding unchosen population, presenting low pre and post-synaptic activity. After reward delivery, the transient elevated activity in the corresponding chosen populations will drive Hebbian plasticity gated by a global RPE (5, 6, 19) (Fig. 1C and D) (see Methods). Thus, only the recurrent synapses of the corresponding chosen populations will modify their synaptic efficacy. Two conceptually different predictions from the neural integrator model can be directly derived from the synaptic model:

1. RPE acts as a global signal gating plasticity in the three-factor learning rule of the recurrent synapses of the value network.
2. Selective feedback from the action selection network is needed for value learning in the valuation network.

The dynamic mechanism for maintaining and updating action values is qualitatively different for the synaptic model compared with the neural integrator. For the synaptic model, action values are maintained in a single fixed-point attractor in neural space. After the reward is collected in the task, a global RPE signal will gate synaptic changes through a three-factor learning rule. Changes in synaptic weights stir the location of the fixed-point attractor in neural space (see Fig. 1C). Therefore, in this model, changes in synaptic weights are needed for action value update.

### Foraging behavior

Can we differentiate these models during foraging in the DFT?

At a single trial level, both models display value-dependent persistent activity during the inter-trial interval (ITI) (Fig. 1E and F; Fig. S1 in SI) as it has been observed experimentally (3, 4). After the action selection network makes a choice, the activity is modified in both models, and action value representations are updated and persist during the next ITI (see Fig. 1D and E) (3, 4). Importantly, as discussed above, in the synaptic model, selective choice signals in the valuation network feedback from the action selection network before reward delivery for action value update. For the neural integrator, these signals are not present by construction (see Fig. 1D vs. E) since they would disrupt action value encoding (see next section).

During a simulated DFT session the reward delivery is stochastic, and the probability of reward for the two contingencies changes in blocks of 100 trials (see Methods), making it necessary for the models to update the value representations to maximize the reward consumption. The instantaneous proportion of choices closely follows the instantaneous proportion of baited reward for the two models (Figs. 2A and B). This dynamical matching is also displayed in a block scale in which, for each block, the slope of the choice ratio, which is the slope of the cumulative choice of contingency A vs. B, matches approximately the baited probability ratio for each block. (2–4, 15) (Figs. 1E and F). We found that the two models display probability matching (14) for a large parameter range.

**Fig. 2.**
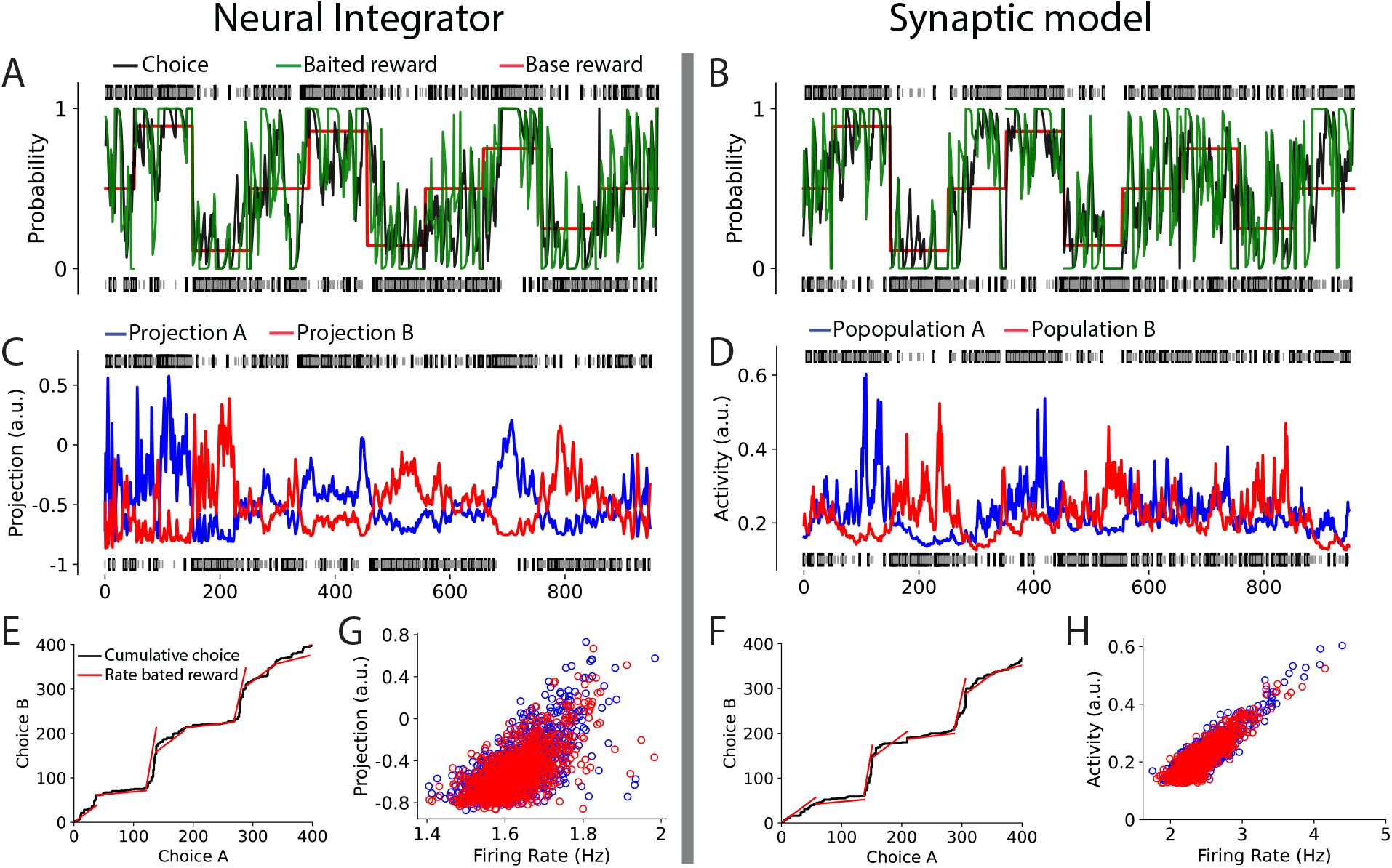
Foraging behavior for the neural integrator (left) and synaptic model (right). (Top row) Foraging behavior during a DFT session (A and B). The base reward probability (red) changes in blocks of 100 trials (see Methods), and the network adjusts its choices, maximizing the baited reward. Black and gray ticks indicate the rewarded and unrewarded trials respectively with choices A (top) and B (bottom). The network’s proportion of choices (black trace) closely follows the proportion of baited reward (green trace) which are smoothed using a causal Gaussian kernel with a standard deviation of 4 trials. (C) Mean projections of the valuation network’s activity on the line attractors axes during the ITI across trials. (D) Mean activity of the valuation network’s selective populations during the ITI across trials. (E and F) Cumulative choice (black) and rate of baited rewards for the neural integrator and synaptic model, respectively. Mean projections on the line attractors axes (G) and mean activity of the selective populations (H) on the valuation network vs. mean firing rate on the action selection network during the ITI.

For the neural integrator, the action value representations encoded in the population activity projected in the line attractor axes dynamically change across trials and they are correlated with the baited rewards (Fig. 2C). Similarly, the synaptic model’s selective populations firing rates during the ITI encoding action value also correlate with the baited reward (Fig. 2D). Overall, both models display qualitatively similar dynamics of value encoding across trials consistent with the value representations observed in the cortex during foraging (3, 4).

The ITI activity in the action selection network also presents, a rather small, action value encoded in its persistent firing. The average firing rate is correlated to the action value encoded by the valuation network in both models (Fig. 2G and H) due to the feedforward connections from the action value network to the valuation network (Fig. 1B and D).

The trial-by-trial dynamics of action value representations both models is mathematically very similar to the Q-learning algorithm in Reinforcement Learning (16) (see Methods and SI). As in the Q-learning algorithm, maintained action values bias action selection performed in our case by the action selection network. Also, as in the Q-learning algorithm, RPE signals are integrated by updating action values in both models. In the neural integrator, this is performed by the neural activity itself, while in the synaptic model, by changes in the synaptic efficacy.

We fit a reduced version of both network models using mouse behavior during the DFT (see Fig. 3, Fig. S2, and the SI for a description of the fitting procedure). We found that both models have comparable performance with standard behavioral models for foraging (3, 4, 20, 21).

**Fig. 3.**
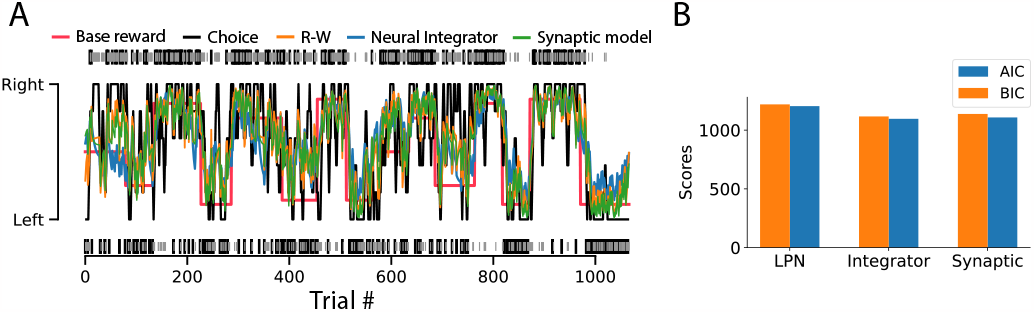
Foraging behavior for a mouse vs. network models during an example DFT session. (A) Foraging behavior of a mouse during a DFT session (see Methods). The base reward probability (red) changes in blocks. Black and gray ticks indicate the rewarded and unrewarded trials respectively for right (top) and left (bottom) choices. Black trace: the proportion of choices smoothed using a running average of 10 trials. The behavior was fitted using a Rescorla–Wagner model (21) (R-W) (orange), the neural integrator (blue), and a synaptic model (green). (B) Model fitting AIC and BIC scores. In this task, the contingencies are licking left and right (see Methods).

Importantly, although both models display qualitatively similar dynamics across trials for encoding value, they have qualitatively different network mechanisms for value encoding and update. These differences can be uncovered by their response to perturbations, as shown in the next section.

### Response to perturbations in neural activity

A central difference between these models is their response to perturbations.

When a perturbation targets action value encoding populations in the valuation network during the ITI the models behave qualitatively differently. In the case of the synaptic model, because action values are stored in the synaptic efficacies, disruptions in the action value encoding due to selective perturbations always rebound to previous values (see Fig. 4A). By contrast, in the case of the neural integrator, perturbations of population activity during the ITI can modify the action value encoding (Fig. 4B). Importantly, however, the effect highly depends on the way of perturbation (Fig. 4D). While on-manifold perturbation dramatically modifies action value encoding, random perturbations have to be much stronger (∼3-5 times than on-manifold perturbations) to have comparable effects to on-manifold perturbations (Fig. 4B and Fig. S3). Uniform (spatially correlated) perturbations where all neurons are perturbed by the same constant amount, reminiscent of optogenetic manipulations, could also disrupt action value encoding as on-manifold perturbations if they are strong enough (∼3-5 times than on-manifold perturbations) (Fig. 4B and Fig. S3). Importantly, strong input noise degrades value encoding leading to drifts in the projections of neural activity in the line attractor axes (Fig. 4C). In the neural integrator, perturbations can be further engineered by ranking neurons by the strength of their encoding to one of the two contingencies. We found that perturbing the first 10% of the highest action value encoding neurons produce an 83% of change in action value encoding (Fig. 4D).

**Fig. 4.**
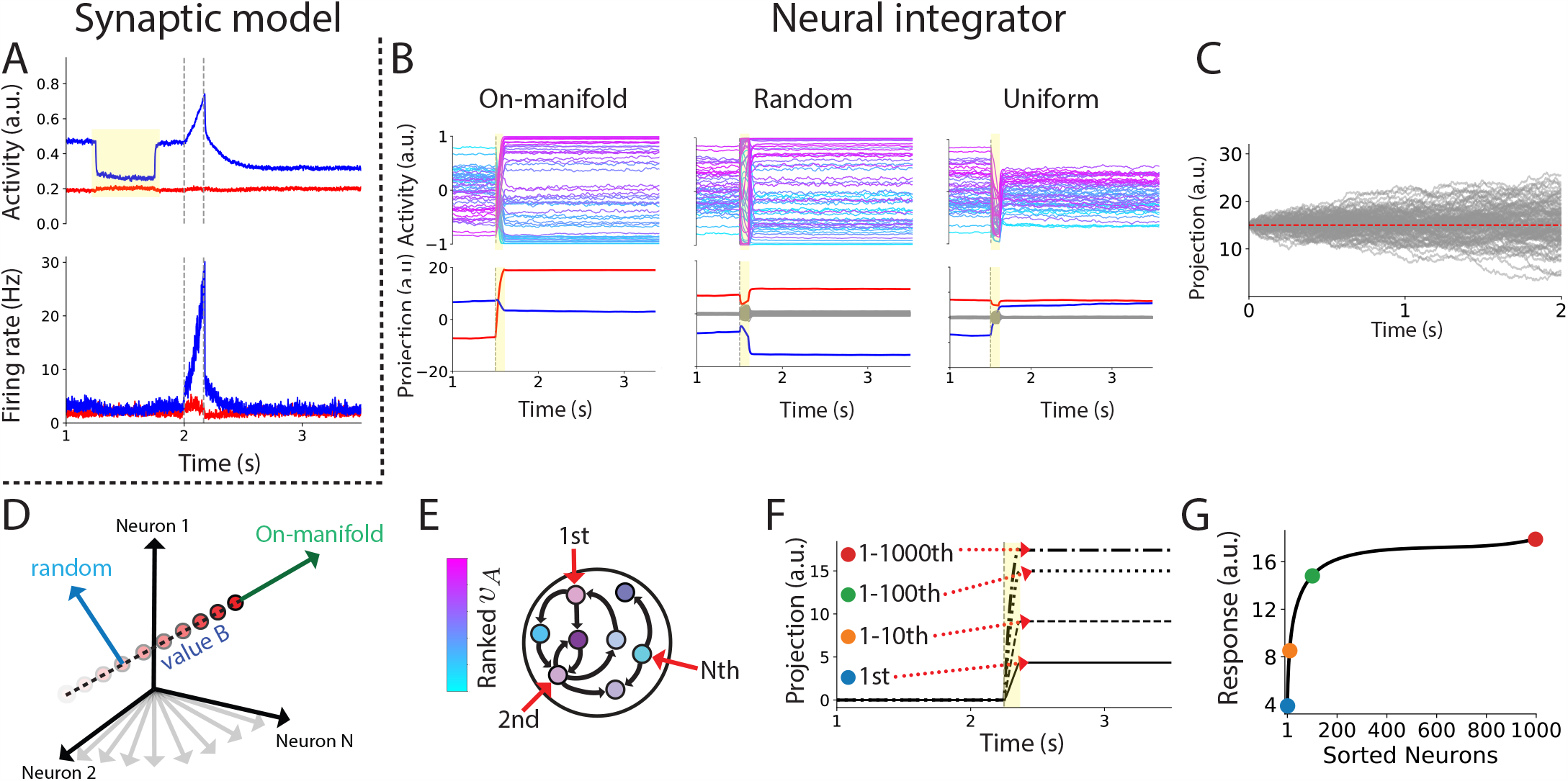
Response to perturbations: neural integrator vs. synaptic model. (A) Top panel: valuation network’s response to a perturbation during the ITI (yellow-shaded region) for the synaptic model. Bottom panel: Action selection network’s response. (B) Examples of on-manifold, random, and uniform brief (100ms) perturbations (starting at the vertical dashed lines and indicated at the yellow-shaded region) in the valuation network for the neural integrator. Top panel: Activity of 50 representative neurons color-coded according to the magnitude of the corresponding entries of the first line attractor axis 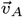 . Random and uniform perturbations are four times larger in magnitude than the on-manifold perturbation. For the stimuli’s parameters see the SI subsection “Three classes of external perturbations”. (C) Projection drift due to noise for 1000 realizations. The dashed red line indicates the initial projection’s value. (D) Schematic of on-manifold and random perturbations. (E) Schematics for the engineered perturbation experiment for the neural integrator. Neurons are ranked according to the entries of the line attractor axis corresponding to one of the contingencies 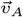 . (F) Change in action value encoding when the 1st, 1-10th, 1-100th, and all ranked neurons are stimulated as in B with the same magnitude. (G) number of stimulated neurons vs. the magnitude of the perturbation response (see SI).

In the synaptic model, the timescale of response to random fluctuations of the selective populations in the valuation network is correlated with action values. In contrast, in the neural integrator, it is not (Fig. 5). The reason is that in the synaptic model, an increase (decrease) in the strength of excitatory recurrent slows down (speeds up) the network’s dynamics (see SI). The neural integrator does not rely on synaptic efficacy changes for action value updates. As a result, there is no association between action value encoding and the time-scale of fluctuations.

**Fig. 5.**
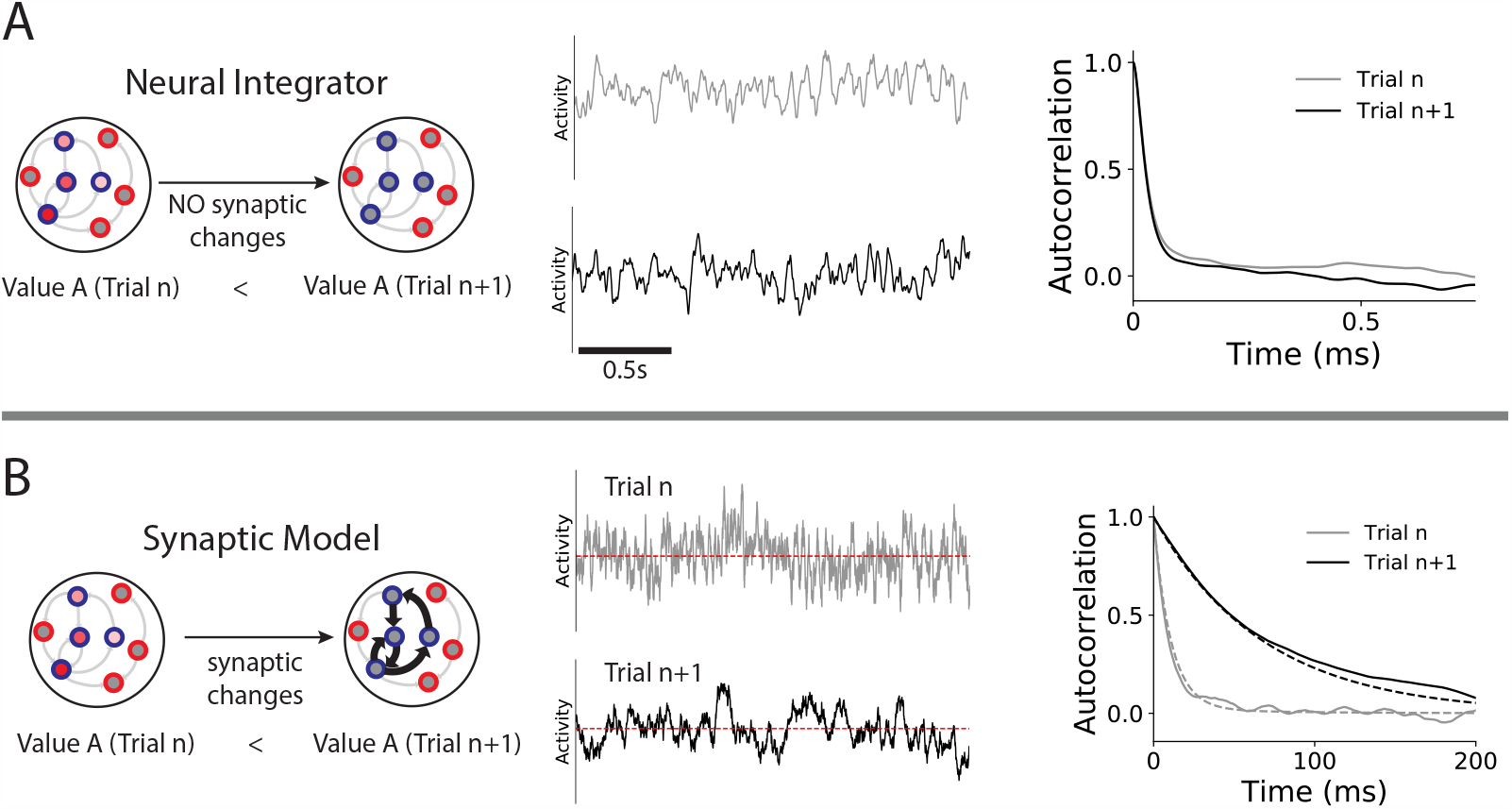
Time scale of the response to fluctuations: neural integrator (A) vs. synaptic model (B). (Left) Schematics of the synaptic changes for two consecutive trials during foraging. (Middle) Network dynamics during the ITI for two consecutive trials. (Right) Autocorrelation function (AF) for two consecutive trials. The AF is also computed analytically for the synaptic model (dashed lines; see SI).

### Response to connectivity perturbations

Synaptic strengths fluctuate in time due to unreliable synapses (22), short-term (23), and long-term plasticity (24). How do synaptic fluctuations affect action value encoding in both models?

In our neural integrator, for constructing the line attractor axes, we set the two eigenvalues corresponding to these axes to have a real part equal to zero (see Fig. 6A and Eq. (28)). These eigenvalues are fine-tuned, and slight departures of a distance *ϵ* from the imaginary axis cause neural activity to drift with a time scale *τ ∝* 1*/ ϵ* (11, 25) (Fig. 6B). Therefore, our model predicts that for slight departures to *ϵ* = 0, action value encoding drifts during the ITI. However, cellular mechanisms can be incorporated into neural integrators to attenuate this drift (25) (see Discussion). When multiple action values are encoded in the neural integrator, the probability of encountering significant drifts in at least one of the line attractor axes due to connectivity perturbations increases with the number of action values being maintained (refer to Fig. S4A and B).

**Fig. 6.**
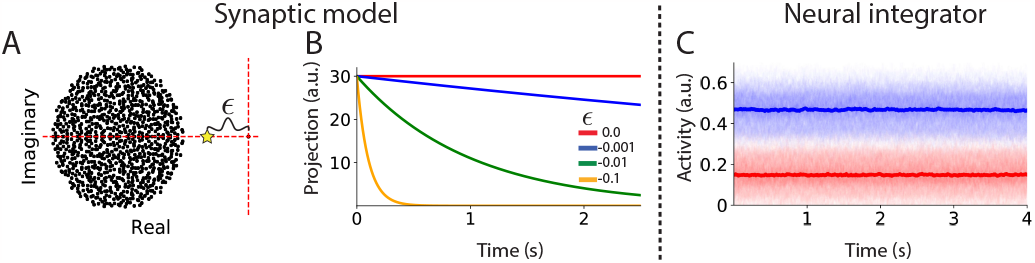
Response to synaptic perturbations: synaptic model vs. neural integrator. (A) Eigenvalue spectra of the neural integrator connectivity **L** (Eq. (29) in Methods). The eigenvalues corresponding to the line attractor axes (Fig. 1A) are at a *E* distance from the imaginary axis (i.e., zero real value). (B) The exponential time decay of the network activity projected on one of the line attractor axes for different values of *ϵ*. (C) The effect of random synaptic perturbations on the synaptic model (see Synaptic perturbations in SI). Each excitatory population A (blue) and B (red) are comprised of *N* = 200 neurons (faint lines). The average activity across neurons for each selective population is shown with tick lines color coded accordingly.

In contrast to the neural integrator, the average neural activity in the synaptic model is robust against synaptic fluctuations. Random synaptic strength fluctuations (see Synaptic perturbations in SI) do not disrupt the average value encoding (Fig. 6C). This is also the case when multiple action values are maintained (see Fig. S4C).

Overall, the synaptic model exhibits significantly greater robustness against synaptic perturbations in the encoding of action values compared to the neural integrator.

### Possible mechanisms for anti-correlated action value updates

In both models, values are independently updated across trials, similar to model free reinforcement learning algorithms (16), in which action values are updated using only the previous history of choices and rewards (see SI, Fitting network models on behavioral data). However, hidden causal structures in the environment can often be used for optimally updating action values during foraging. For example, this is the case in variations of the DFT in which the changes in the probability of rewards are strongly anti-correlated (26). In this case, opposite action values updates are optimal for maximizing reward during the task. What are the circuit mechanisms for anti-correlated action values updates in structured environments? We explored two possible scenarios in the neural integrator and the synaptic model.

For the neural integrator, one possible scenario for anticorrelated action value updates is that the line attractor axes are anti-correlated (Fig. 7A left), and a contingency-dependent RPE signal is aligned with one of the axes. In this case, RPE signals produce anti-correlated action values updates (Fig. 7A right). An alternative scenario is that the RPE signal is comprised of two inputs, each one aligned with one of the two line attractor axes and with opposite signs (Fig. 7B left). This scenario also leads to opposite changes in action value representations (Fig. 7B right).

**Fig. 7.**
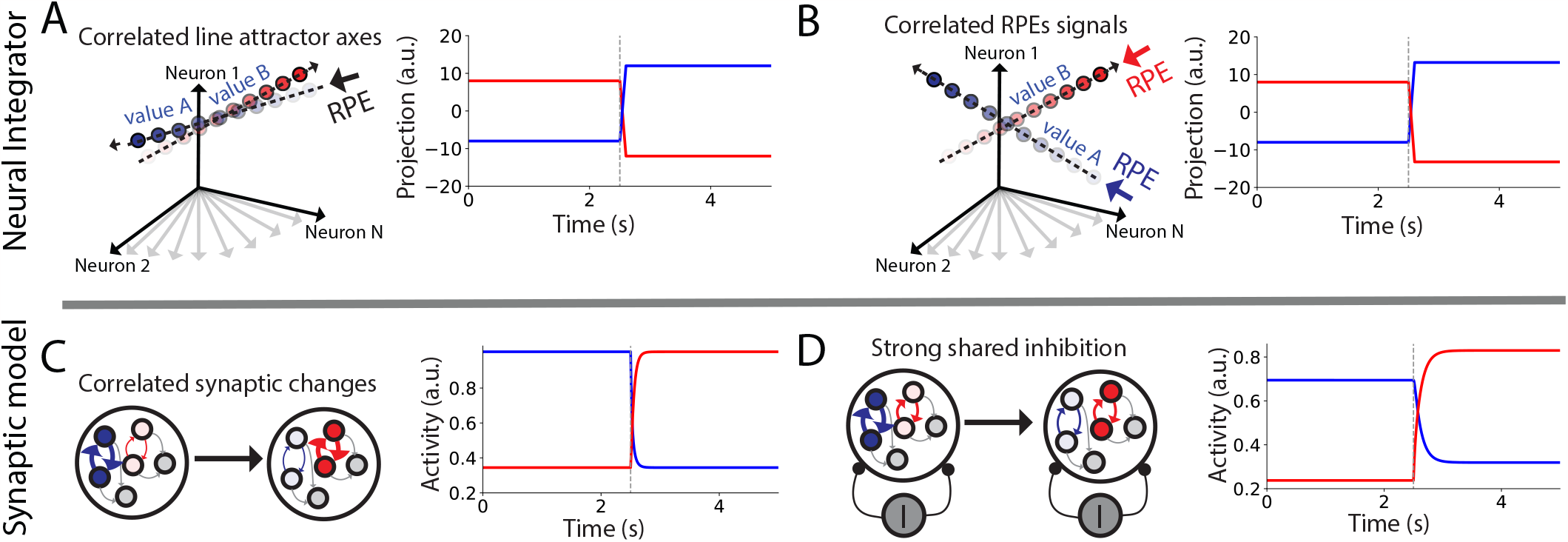
Possible mechanisms for anti-correlated action value updates. (A) Left: Action values encoded in anti-correlated line attractor axes are updated in opposite directions by an RPE signal aligned to one of the axes. (B) Left: Action value encoding update RPE signal comprised of two inputs, each one aligned with one of the line attractor axes but with opposite signs. Right (A and B): Projections dynamics. (C) Left: Anti-correlated synaptic changes in the synaptic model. (D) Left: Synaptic changes in the synaptic model of only one selective population produce anti-correlated action value updates due to strong inhibition. Right (C and D): Population activity dynamics. The RPE signal is delivered and the synaptic update is performed at 2.5s (dashed line). The RPE signal has a duration of 100ms.

For the synaptic model, anti-correlated synaptic changes (Fig. 7C left) for the corresponding selective populations produce anti-correlated action value updates (Fig. 7A right). An alternative scenario relies on strong shared inhibition (Fig. 7D left) that anti-correlates neural activity (see SI, synaptic model). In this scenario, synaptic changes in only one selective population produce anti-correlated action value updates (see Fig. 7D right and SI).

## Discussion

We investigated two contrasting models for action value encoding and update mechanisms in foraging behavior: a synaptic model and a neural integrator. The synaptic model maintains action values in synaptic strength and updates them through synaptic plasticity, whereas the neural integrator maintains action values in a continuous attractor through neural activity and updates via on-manifold perturbations. It is worth noting that we use an extension of the concept of line attractor (11) to several line attractors in a high dimensional state space of neural population activity (12).

Our models successfully replicate both choice behavior and key observations in neuronal recordings. Specifically, our modeling requires that RPE signals in the neural integrator are vectorial and action-specific on-manifold signals, whereas the synaptic model exhibits global RPE signals. Furthermore, we found that the neural integrator is more susceptible to perturbations in neural activity and connectivity, while the synaptic model displays remarkable robustness against such perturbations.

When examining anti-correlated activity updates necessary for optimal performance in foraging tasks with hidden causal structures, we found that the synaptic model efficiently accommodates these updates by solely increasing the amount of shared inhibition, offering a parsimonious explanation for the synaptic changes necessary for learning new tasks structures Our initial findings suggest that the synaptic model is a compelling candidate for the maintenance and update of action values in the brain, given its robustness and because its observed dynamics align with multiple experimental observations. We identified requirements for neural integrator model that presently lack experimental support, including the assumption that different dopamine neural populations are selective for action values and project to a valuation system in such a way to be well aligned with the line attractor axes. Overall, our work provides a computational framework to guide future research in order to disprove these two candidate mechanisms for foraging in the brain.

### Network architecture

Consistent with the observed functional separation between valuation and action selection in the cortical-basal ganglia-thalamocortical limbic and motor loops during decision-making (13, 17), we build a network model with two distinct connected networks that separately realize valuation and action selection. In the action selection network, actions are selected by a winner-take-all dynamics elicited by a transient go signal. Importantly, in our models (both the synaptic model and neural integrator), this go signal acts in a non-selective multiplicative fashion transiently modifying the overall recurrent connections of the action selection network (see Eq. (8-12) on Methods) and momentarily creating two stable attractors that represent chosen actions (27, 28) without the need of reset the network. This multiplicative mechanism is consistent with experimental evidence in decision-making tasks in which go signals input the thalamus from the midbrain (29). These inputs can effectively increase the recurrent connectivity in cortex due to the recurrent thalamocortical loop (30, 31).

Due to the feedforward connections from the valuation network to the action selection network, the ITI activity in the action selection network also presents a rather small, action value encoding in its persistent firing (Fig. 1E and F; Fig. 2G and H). Our models’ predictions are consistent with the observed value encoding in the rat’s secondary motor cortex during foraging (32).

Our models provide contrasting predictions for the feedback projections from the action selection to the valuation network. In the neural integrator, there is no feedback from the action selection network. This aims to avoid choice signals disrupting action value encoding. It is expected that, in this scenario, any feedback projections from the action selection network to be weak or approximately orthogonal to the line attractor axes, to maintain the action value encoding. In contrast, in the synaptic model, choice signals feedback from the action selection network are necessary for reward-dependent learning (see Methods). In this scenario, we expect feedback from the action selection to the valuation network to play a causal role in learning.

### Synaptic model

In the valuation network of our synaptic model, recurrent connections undergo reward-modulated plasticity. Trial-to-trial changes in its recurrent synaptic strengths cause a graded variation of neural activity during ITIs in proportion to action values updates that are consistent with recorded activity in the mPFC and RSC (3, 4). Our results contrast with classic synaptic models for action value maintenance and update during foraging in which plasticity is limited to the feedforward input projections (5, 6). In these models, action selection is biased by action value encoded in the strengths of these feedforward projections and the network performs action selection by a winner-take-all dynamics that resets after each trial, at odds with recent experimental observations (3, 4).

Recent work studying foraging on the fly suggests that synaptic plasticity might be the mechanism underlying foraging behavior in this animal (33).

In the synaptic model, the timescale of response to random fluctuations of the selective populations is correlated with action values. Consistent with this prediction, such dependence on the autocorrelation function with value has been observed in value-based tasks in monkeys (34).

### Neural integrator

For the neural integrator model, we assumed two line attractors encoding action values (see Fig. 1A). We showed that this model can account for graded changes of neural firing during ITIs, but requires that value updating is realized by an action-specific vectorial RPE signal projecting to value-coding neurons in a special way along a specific action value axis (e.g., for action A, not B), in a given trial. These signals could be mediated by dopamine cells (35), norepinephrine cells (36), or subcortical input (37). The latter implies that these projections target selective neural populations in a choice-dependent manner, an assumption not supported in the case of dopamine by presently known evidence but testable in the future.

### Robustness to perturbations

In the synaptic model, the dynamics during the ITI is given by a single stable fixed-point attractor. We predict that optogenetic perturbations in the neural activity during the ITI recover to similar firing rate levels, enabling the robust encoding of action value against perturbations.

For the line attractor, the effect of a perturbation in neural activity diminishes as it becomes more misaligned with the line attractor axes. However, our findings indicate that even a strong perturbation with minimal alignment to the line attractor axes can result in significant disruptions to action value encoding. We predict that brief and strong optogenetic perturbations during the ITI will disrupt action value encoding in this scenario. We also predict that optogenetic perturbations can be engineered to maximize its effect by targeting the neurons with higher action value encoding. Random persistent fluctuations induce drift in action value encoding, predicting deterioration of the action value maintenance for long ITIs. Our study primarily focuses on linear neural integrators or those with weak non-linearities. It is possible that incorporating strong nonlinear effects (38) or the implementation of an approximately continuous line attractor consisting of multiple discrete stable attractors (39) could produce some enhancement in the robustness of the network.

In the neural integrator, the connectivity is fine-tuned, resulting in a manifold of marginally stable states. Even small perturbations in the synaptic connectivity caused by fluctuations in the synaptic efficacy can lead to persistent drifts in neural activity. Furthermore, as the number of action values maintained increases, the likelihood of significant drifts in the encoding of action values also rises. Consequently, the neural integrator is very susceptible to synaptic perturbations. In contrast, the synaptic model exhibits stability against synaptic fluctuations. Action values in this model remain largely unaffected by such perturbations.

Overall, our findings show that the synaptic model provides a significantly more robust mechanism for action value encoding.

### Structured tasks

In both models, action values are updated depending on the history of choices and rewards, with no use of the specific structure of the task. Furthermore, our network models are algorithmically very similar to the Q-learning algorithm (16) (see SI). Consequently, from a reinforcement learning standpoint, our models are considered *model free* (16), which refers to the fact that they do not use information about the environment’s structure for performing the action values updates encoded in the synapses and neural activity in the respective models.

In many situations, it is advantageous to learn the causal structure of the world for maximizing reward consumption (40, 41). During value-based decision-making in uncertain environments with underlying hidden structures, inferring the causal structure of the task can lead to higher rewards (26, 42–44). One interesting scenario for foraging is a recently proposed task in which a hidden state induces anti-correlated and extreme changes in reward probabilities (26).

In our models, anti-correlated updates of the firing rate activity of selective populations could capture the hidden structure of this task. In the neural integrator, anti-correlated line attractor axes lead to anti-correlated action-value updates. However, adapting to new task structures may involve learning new alignments between line attractor axes effectively changing the manifold structure, which may require global changes in synaptic connectivity. It is unclear whether this can be flexibly achieved using local learning rules. In contrast, anti-correlated updates can also be achieved in the line attractor by adjusting the RPE alignment with the line attractor axes. Consequently, when learning new tasks, only modifying RPE signal alignments becomes necessary in this scenario. To our knowledge, there is no evidence of such RPE characteristics in the dopamine system.

In the synaptic model, anti-correlated changes in firing rates and action value encoding can be achieved using local three-factor learning rules by leveraging the amount of shared inhibition in the network. Our results demonstrate that if there is strong shared inhibition among selective populations, synaptic changes in the recurrent connections of one population give rise to anti-correlated changes in firing rates. Interestingly, we hypothesize that for learning new tasks inhibitory plasticity (45, 46) could sculpt the correlation structure of the firing rate updates. This leads to a possible flexible mechanism for learning new causal structures in the environment.

### Foraging in naturalistic environments

While animals are foraging in a naturalistic environment, they must remember the expected reward at each location and estimate the travel cost to other locations to decide on actions (47). In our model, we consider an environment in which there is only one location and two action options that are simultaneously presented. By contrast, in more naturalistic foraging, different choice alternatives (e.g., food patches) are present at different time points (48–50). For example, if a limited and fixed number of actions are taken in each state, e.g., stay vs. leave, if leave goes left vs. right. It would suffice to maintain in memory the location values by a valuation network as shown in Fig. S4 in the SI. An action selection network model similar to the one presented in this work can decide whether to stay or leave and the direction of travel by comparing the value of the current location (state) vs. the integrated value of the rest of the environment discounting travel costs. It remains to be seen how our model can be extended for such foraging in naturalistic environments.

## Materials and Methods

### Foraging task

#### Dynamic foraging task

We implement a dynamic foraging task (DFT) similar to the one in (3, 4) in both our network simulations and behavioral experiments. Briefly, in this task, after a go cue (auditory for the mice), the subject chooses freely among two alternatives (by licking one of two custom-built lick ports for the mice) that deliver a reward (water for the mice) probabilistically. After an inter-trial interval (ITI) of the order of several seconds (3s in the case of the network and 5.79s median ITI in the case of the mice), the next go cue is presented. Depending on their choice, a reward is delivered with a probability that changes randomly in blocks of 100 trials for the network (see Fig. 2A and B). For the behavioral experiments in mice, the number of trials in each block is drawn from a bounded exponential distribution ranging from 40 − 100 trials. For this task, the reward schedule uses baited rewards (e.g., see (2–4, 20)), that is that once the reward is assigned, it remains in the corresponding contingency (network)/port (mice) for its consumption. The reward probabilities for the two contingencies (*p*_*A*_, *p*_*B*_) (network)/(*p*_Right_, *p*_Left_) (mice) were chosen randomly from {(0.225, 0.225), (0.4, 0.05), (0.05, 0.4), (0.3857, 0.0643), (0.0643, 0.3857)*}*.

#### Animal procedure and behavioral task

All animal procedures were in accordance with protocols approved by the Janelia Research Campus Institutional Animal Care and Use Committee and have been described before (51, 52). Briefly, C57BL/6J mice were each implanted with a titanium head post (dx.doi.org/10.17504/protocols.io.9a8h2hw) and housed individually in a reverse 12-hour dark/12-hour light cycle. After recovery from the head post surgery, the mice were water restricted and then habituated for 2 days prior to task training.

Behavioral sessions lasted 1-2 hours during the dark phase. Mice were head restrained while resting in a 31.8 mm acrylic tube (8486K331, McMaster-Carr) inside a dark and sound-attenuated box. The go cue (3 kHz, 100 ms) was presented by a speaker (TW025A20, Madisound). Water reward (2-4 μL) was delivered by solenoid valves (LHDA1233215H, The Lee Co) through two lick ports, spaced 4.5 mm apart (8987K47, McMaster-Carr). Licks were detected by a custom circuit (JF-SV-LP0001, Janelia Research Campus). Task events were controlled by a Bpod State Machine (Sanworks) and programmed in PyBpod (https://github.com/hanhou/Foraging-Pybpod).

## Network models

### Network architecture

As described in the main text, we develop two network models for contrasting two different mechanisms for action value maintenance and update during foraging: (i) neural integration and (ii) synaptic learning. Both models have similar network architecture consisting of a mesoscopic network of two inter-connected areas. Each area is modeled as a recurrent network. Action values are maintained and updated depending on reward in a network we refer to as the valuation network (see Fig. 1B and D). For action selection, action values are read out from the valuation network and projected for biasing actions in a network we refer to as the action selection network (see Fig. 1B and D). The dynamics of the action selection network is the same for both models.

As described below, two critical differences exist between the neural integrator and the synaptic model architectures. First, there are feedback projections between the action selection and the valuation network in the synaptic model, while in the neural integrator, these projections do not exist. Second, in the neural integrator, reward prediction error (RPE) signals are selectively projected to the two action value axes, while in the synaptic model, RPE is a global signal projected to the whole valuation network (see more in sections below).

### Action selection network

For the action selection network, we use a reduced *mean field model* that has been shown to be a good approximation of the dynamics of a full recurrent network of spiking neurons (28). This reduced model represents the dynamics of the fraction of activated N-methyl-D-aspartate (NMDA) receptors of two selective excitatory populations. These populations represent the two contingencies. The dynamics is described by the below equations

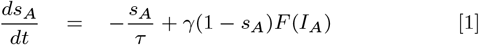

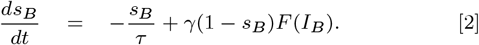

with *γ* = 0.641 and *τ* = 100ms. The function *F* is the input-output transfer function given by

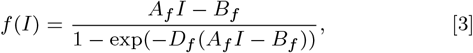

where *A*_*f*_ = 270Hz/nA, *B*_*f*_ = 108Hz, and *D*_*f*_ = 0.154s (28, 53).

The synaptic currents are given by a recurrent (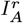 and 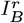), a background synaptic noise (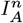 and 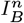), and a non-selective external go signal (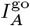 and 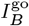) components

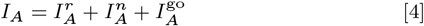

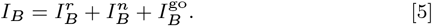

The recurrent component is given by

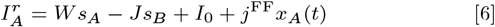

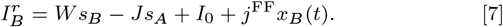

Here *W* = 0.2609 corresponds to the average recurrent synaptic weights for each selective population, *J* = 0.0497 corresponds to the effective shared inhibition weights, and *I*_0_ = 0.3381 to the background current. For these parameter values, the action selection network is in a winner-take-all regime suitable for decision-making (28). The parameter *j*^FF^ corresponds to the feedforward connections from the valuation to the action selection network. The variables *x*_*A*_(*t*) and *x*_*B*_(*t*) represent the mean activity of the two selective populations *r*_*A*_(*t*) and *r*_*B*_(*t*) in the case of the synaptic model and the two projections *m*_*A*_(*t*) and *m*_*B*_(*t*) in the case of the neural integrator respectively (see following sections for the detailed implementation of the valuation network). For the synaptic model *j*^FF^ = 0.2 for Fig. 1F and *j*^FF^ = 0.04 for Fig. 2B, D, F, and H. For the neural integrator *j*^FF^ = 0.0015 for Fig. 1B and *j*^FF^ = 0.003 for Fig. 2A, C, E, and G. The equations for the background synaptic noise due to AMPA synapses are given by

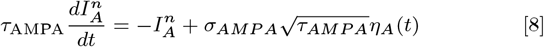

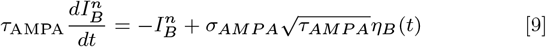

Here *η*_*A*_(*t*) and *η*_*B*_(*t*) are Gaussian noise variables. The time scale of AMPA synapses is *τ*_AMPA_ = 2ms. Lastly, the intensity of the noise for the synaptic model is *σ*_*AMP A*_ = 0.003 for Fig. 1 and Fig. 4 and *σ*_*AMP A*_ = 0.03 for Fig. 2. For the neural integrator is *σ*_*AMP A*_ = 0.003 for Fig. 1 and *σ*_*AMP A*_ = 0.02 for Fig. 2.

In our model, the go input currents 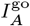 and 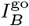 correspond to transient non-selective inputs that lasts un^*A*^til one o^*B*^f the populations reaches a set firing rate (*F* (*I*_*A*_) or *F* (*I*_*B*_)) threshold of 30Hz. Consistent with recordings in the Anterior Lateral Motor cortex (ALM) during motor initiation (54) and as proposed in theoretical studies (30, 31), we hypothesize transient non-selective go signals gate actions to the motor thalamus. Since the recurrent thalamocortical loop involving ALM and ventromedial (VM) thalamic nucleus is causally involved in motor initiation (29, 55), the net effect of an input to the thalamus corresponds to transient effective synaptic weights (30, 31) given by

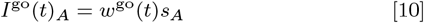

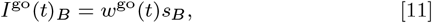

where

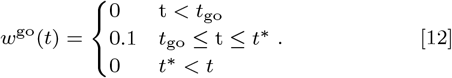

We define *t*^∗^ as the time when one of the populations firing rates *F* (*I*_*A*_) or *F* (*I*_*B*_) reaches the threshold of 30Hz.

The numerical integration time step used was *dt* = 0.1ms.

### Network dynamics

For the synaptic model, the network dynamics are given by the linearized dynamics of a network of two excitatory selective populations to the two contingencies in the DFT with shared inhibition

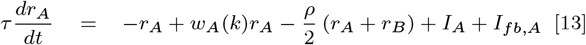

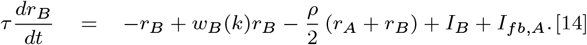

Here the variables *r*_*A*_ and *r*_*B*_ correspond to the mean firing rates of the two selective populations. The parameters *w*_*A*_(*k*) and *w*_*B*_(*k*) correspond to the mean synaptic efficacy of the recurrent connections in the trial *k*. The synaptic efficacies are plastic and change on a trial-to-trial basis according to a learning rule specified in the next section. The parameter *ρ* = 0.05 corresponds to the effective, shared inhibition. The background input current is given by the parameters *I*_*A*_ = *I*_*B*_ = 0.1. The timescale of the dynamics is taken as *τ* = 10ms. The numerical integration time-step used was *dt* = 10ms. The currents *I*_*fb,A*_ = *w*_*fb*_*s*_*A*_ and *I*_*fb,B*_ = *w*_*fb*_*s*_*B*_ correspond to the feedback current from the action selection network. We use *w*_*fb*_ = 0.2. Since during the ITI the feedback current is small, for the below analysis, we will assume *I*_*fb,A*_ = *I*_*fb,B*_ = 0. In Fig. 7C *w*_*A*_ = 0.87, *w*_*B*_ = 0.62, and *ρ* = 0.25. In Fig. 7D *w*_*A*_ = 0.87, *w*_*B*_ = 0.62, and *ρ* = 0.45.

Although extremely simple, this model captures essential features of an E-I network and has the advantage that several quantities can be calculated analytically. This model has a single fixed-point attractor corresponding to the dynamics’ stationary state. We analytically calculate its fixed point (see section Synaptic model on SI).

### Synaptic model’s reward-dependent learning rule

In the synaptic model, the recurrent connections synaptic efficacies of the two selective populations in the valuation network *w*_*A*_(*k*) and *w*_*B*_(*k*) are learned using a reward-dependent plasticity rule (5, 6, 19) in a trial-by-trial (where *k* is the trial index) basis. In our three-factor learning rule the synaptic changes are given by

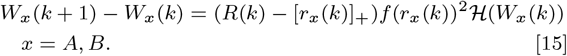

Here the reward is a binary variable *R* ∈ {0, 1*}*. The function [*•*]_+_ is the rectifier linear function with a minimum equal to 0 and a maximum equal to 1. The function *ℋ* (*x*) given by

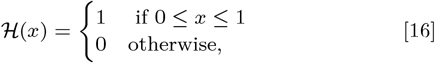

enforces a hard constraint on the synaptic weights preventing a firing rate instability. This could be implemented by homeostatic plasticity mechanisms (56).

The reward prediction error (RPE) at trial *k* is given by the difference RPE(*k*) = *R*(*k*) − [*r*_*x*_(*k*)]_+_. Here *r*_*x*_(*k*) is the mean firing rate of the population in the valuation network corresponding to the chosen contingency *x* = *A, B* at trial *k*. Then Eq. (15) reads as

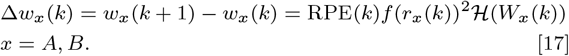

We assume that the RPE is computed elsewhere in the brain, for example, in the Ventral Tegmental Area (VTA) (18).

The Hebbian term of the three-factor learning rule (19), which is the product of a non-linear function of pre and post-synaptic activity of the selective populations, is given by the term *f* (*r*_*x*_)^2^ = *f* (*r*_*x*_) · *f* (*r*_*x*_) for *x* = *A, B*. Notice that the Hebbian term is squared because the pre and post-synaptic activity is the activity of the selective populations *A* or *B*. This term will modify the recurrent connections of the selective populations *w*_*A*_(*k*) and *w*_*B*_(*k*). In our model, the function *f* is a highly non-linear step function

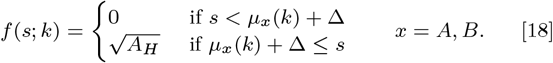

Here *μ*_*A*_(*k*) and *μ*_*B*_(*k*) correspond to the running temporal firing rate average of population *A* and *B* in the valuation network, respectively, at trial *k*. The above function induces plasticity in synapses where the firing rates are ∆ larger than the average firing rate of the previous trial. This class of non-linear plasticity rule that potentiates firing rate outliers has been recently inferred from *in vivo* data (57, 58).

In our model, the running average *μ*_*A*_(*k*) and *μ*_*B*_(*k*) we assume a slow exponential kernel of the order of seconds. Since, in our task, the ITI period is much longer than the response period, we can replace *μ*_*A*_(*k*) and *μ*_*B*_(*k*) by the mean firing rates during the ITI. After the go signal, when the choice is selected by the action selection network, the chosen selective population in the action selection network will elevate its firing rate (see Fig. 1F). The feedback projections from the action selection to the valuation network will produce a transient elevated activity in the corresponding population. In contrast, little to no feedback activity is produced in the corresponding unchosen population (see Fig. 1B, bottom row left). This selective feedback activity in the valuation network and our learning rule in Eq. (18) produce changes in the recurrent synaptic efficacies of only the chosen populations in the valuation network. Therefore, the final learning rule we use in our numerical simulations is

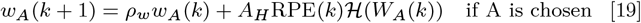

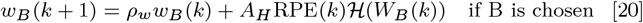

and

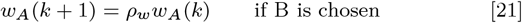

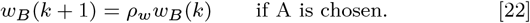

The parameter *ρ*_*w*_∈ [0, 1] represents to a trial-by-trial forgetting parameter. If *ρ*_*w*_ = 1, there is no forgetting across trials, while if *ρ*_*w*_ = 0, the synaptic efficacies modifications are forgotten in one trial. We choose no forgetting *ρ*_*w*_ = 1 for simulations in Fig. 2, and we fit this parameter using behavioral data in Fig. 3 (see the section Fitting network models in behavioral data in SI). The parameter *A*_*H*_ in Eqs. (18-20) corresponds to the maximum amplitude of the Hebbian term in our three-factor learning rule. For the network simulations in Fig. 2 *A*_*H*_ = 0.12 while we fit this parameter using behavioral data in Fig. 3 (see the section Fitting network models in behavioral data in SI for the parameter values of the fit and Fig. S2).

For the synaptic update in Fig. 7C, ∆*w*_*A*_ = − 0.25 and ∆*w*_*B*_ = 0.25 while for Fig. 7D ∆*w*_*A*_ = 0 and ∆*w*_*B*_ = 0.33.

### High-dimensional neural integrator

For constructing a neural integrator in high dimensional neural space, our starting point is a recurrent linear network with *N* number of units. The following linear equations give the dynamics

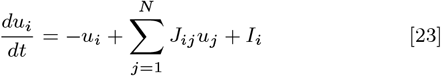

of the synaptic current *u*_*i*_ for each neuron (59). In this network model, the connectivity is random. Each entry is drawn from a normal distribution with zero mean and variance equal to *g*^2^*/N* . The external input to the network is given by the vector 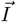;. For all the simulations in the paper, we used *g* = 0.5.

By defining the below matrix

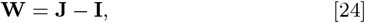

where **I** is the identity matrix. Then Eq. (23) can be written in vectorial form as

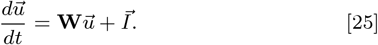

Using the singular value decomposition (SVD) the matrix **W** can be written as the sum of rank-1 matrices scaled by the corresponding singular vectors

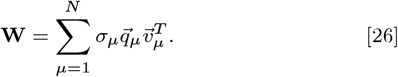

Here the singular vectors are ordered from larger (*σ*_1_) to smaller (*σ*_*N*_). We constructed a neural integrator for encoding two action values similarly as in (12). For constructing a neural integrator with two integration modes, we subtract to the matrix **W** the rank-1 matrices in Eq. (26) corresponding to the two smaller singular values, i.e.,

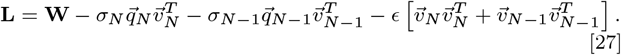

Notice that here introduce a rank-2 normal perturbation that will lead to drifts of time scale *τ* ∼1*/ϵ* in the directions 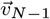 and 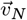 . We then use this new matrix **L** as the connectivity matrix of our neural integrator, which now has two integration modes.

This model can be extended for maintaining *p* values of actions, by constructing a network with *p* line attractor axes:

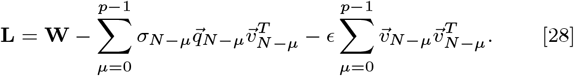

The dynamics of the neural integrator is given by

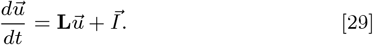

We compute the firing rates as a non-linear transformation of the synaptic currents using a sigmoidal transfer function 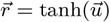. By projecting the network activity 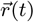 onto the right singular vectors we decompose the activity in *N* network modes

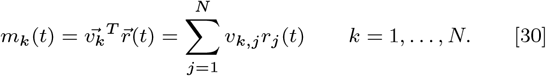

When we project Eq. (29) in the two line attractor axes that maintain the corresponding action values in our DFT we obtain

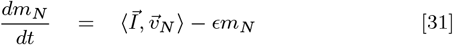

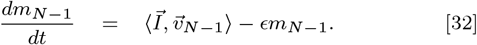

For *ϵ*= 0 reads

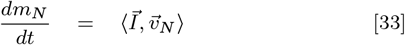

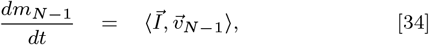

which correspond to the equations of two independent one-dimensional neural integrators. Where ⟨·, · ⟩ corresponds to the inner product of two vectors.

In our model we assume **J** to be independently and identically distributed (i.i.d.) from a Gaussian distribution, i.e., 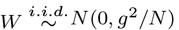. The input current to the network is given by

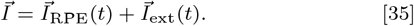

Here 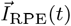 is an input proportional to the RPE that updates the action value after reward delivery by briefly pushing or pulling in the direction of the action value axis corresponding to the chosen contingency (i.e., in the directions 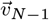 and 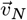 for the contingency A and B respectively). The 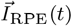 is given by

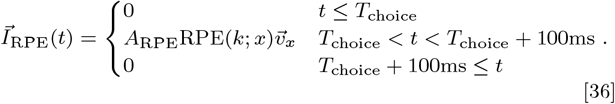

Here the RPE is given by

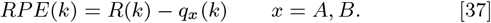

Here *q*_*x*_(*k*) corresponds to the quantile of the projection of the population activity on the line attractor axis corresponding to the chosen contingency (i.e., A or B) (see section Neural integrator on SI). The rationale for using the quantile is to normalize this value to take values between 0 and 1 to be comparable to the reward (i.e., *R* ∈{0, 1}) since the projection can have arbitrary values. We believe this normalization necessary for computing the RPE in our model could be performed by subcortical structures as the VTA (35, 60).

The parameter *A*_RPE_ is the strength by which the RPE signal modifies the population activity. For Fig. 1E *A*_RPE_ = 0.1 and for Fig. 2 *A*_RPE_ = 0.3. The RPE signal stimulates the network after the reward is delivered, which happens instantly after the choice is made at time *T*_choice_ in our model. *T*_choice_ is *T*_*go*_ = 3s plus the reaction time of the action selection network at a given trial and therefore varies from trial to trial.

The 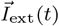 is an external perturbation to the network. For Figs. 1-3 we set 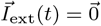.

In Fig.7A and B, a 100ms stimulus is delivered starting at 2.5s. In Fig.7A, the two line attractor axes exhibit perfect anti-correlation, and the stimulus is proportional to one of the line attractor axes, having a norm equal to 2. In Fig. 7B, the stimulus is proportional to 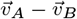, with a norm equal to 3.

The numerical integration time-step used was *dt* = 10ms.

## Software and data availability

The network simulations were performed using custom Python scripts available at the GitHub repository https://github.com/ulisespereira/foraging-integrator-vs-synaptic. The code for fitting the reduced network models on mice behavior is available upon request.

## ACKNOWLEDGMENTS

We thank Nai-Wen Tien and Marton Rozsa for their help with the behavioral training in the pilot experiments. We thank Aldo Battista and Jorge Jaramillo for their constructive comments on the manuscript. This work was supported by The Swartz Foundation (to U.P.-O.), James Simons Foundation Grant 543057SPI (to X.-J.W.), National Institutes of Health grant R01MH062349 (to X.-J.W.), Janelia Visiting Scientists Program (to X.-J.W. and U.P.-O.), Howard Hughes Medical Institute (to K.S.), National Institute of Health U-19 grant 1U19NS123714-01 (to H.H. and K.S.), the Simons Collaboration on the Global Brain (to H.H., K.S., and X.-J.W.), and The Allen Institute for Neural Dynamics (to H.H. and K.S.).

## References

1. ML Platt, PW Glimcher, Neural correlates of decision variables in parietal cortex. Nature. 400, 233–238 (1999).

2. LP Sugrue, GS Corrado, WT Newsome, Matching behavior and the representation of value in the parietal cortex. science 304, 1782–1787 (2004).

3. BA Bari, et al., Stable representations of decision variables for flexible behavior. Neuron 103, 922–933 (2019).

4. R Hattori, B Danskin, Z Babic, N Mlynaryk, T Komiyama, Area-specificity and plasticity of history-dependent value coding during learning. Cell 177, 1858–1872 (2019).

5. A Soltani, XJ Wang, A biophysically based neural model of matching law behavior: melioration by stochastic synapses. J. Neurosci. 26, 3731–3744 (2006).

6. Y Loewenstein, HS Seung, Operant matching is a generic outcome of synaptic plasticity based on the covariance between reward and neural activity. Proc. Natl. Acad. Sci. 103, 15224–15229 (2006).

7. N Frémaux, H Sprekeler, W Gerstner, Functional requirements for reward-modulated spike-timing-dependent plasticity. J. Neurosci. 30, 13326–13337 (2010).

8. JX Wang, et al., Prefrontal cortex as a meta-reinforcement learning system. Nat. neuroscience 21, 860–868 (2018).

9. S Hochreiter, J Schmidhuber, Long short-term memory. Neural computation 9, 1735–1780 (1997).

10. K Krishnamurthy, T Can, DJ Schwab, Theory of gating in recurrent neural networks. Phys. Rev. X 12, 011011 (2022).

11. HS Seung, How the brain keeps the eyes still. Proc. Natl. Acad. Sci. 93, 13339–13344 (1996).

12. S Druckmann, DB Chklovskii, Neuronal circuits underlying persistent representations despite time varying activity. Curr. Biol. 22, 2095–2103 (2012).

13. J Cox, IB Witten, Striatal circuits for reward learning and decision-making. Nat. Rev. Neurosci. 20, 482–494 (2019).

14. RJ Herrnstein, Relative and absolute strength of response as a function of frequency of reinforcement. J. experimental analysis behavior 4, 267 (1961).

15. B Lau, PW Glimcher, Dynamic response-by-response models of matching behavior in rhesus monkeys. J. experimental analysis behavior 84, 555–579 (2005).

16. RS Sutton, AG Barto, Reinforcement learning: An introduction. (MIT press), (2018).

17. H Tang, VD Costa, R Bartolo, BB Averbeck, Differential coding of goals and actions in ventral and dorsal corticostriatal circuits during goal-directed behavior. Cell reports 38 (2022).

18. W Schultz, P Dayan, PR Montague, A neural substrate of prediction and reward. Science 275, 1593–1599 (1997).

19. N Frémaux, W Gerstner, Neuromodulated spike-timing-dependent plasticity, and theory of three-factor learning rules. Front. neural circuits 9, 85 (2016).

20. GS Corrado, LP Sugrue, HS Seung, WT Newsome, Linear-nonlinear-poisson models of primate choice dynamics. J. experimental analysis behavior 84, 581–617 (2005).

21. RA Rescorla, A theory of pavlovian conditioning: Variations in the effectiveness of reinforcement and non-reinforcement. Class. conditioning, Curr. research theory 2, 64–69 (1972).

22. LE Dobrunz, CF Stevens, Heterogeneity of release probability, facilitation, and depletion at central synapses. Neuron 18, 995–1008 (1997).

23. MV Tsodyks, H Markram, The neural code between neocortical pyramidal neurons depends on neurotransmitter release probability. Proc. national academy sciences 94, 719–723 (1997).

24. PJ Sjöström, GG Turrigiano, SB Nelson, Rate, timing, and cooperativity jointly determine cortical synaptic plasticity. Neuron 32, 1149–1164 (2001).

25. S Lim, MS Goldman, Balanced cortical microcircuitry for maintaining information in working memory. Nat. neuroscience 16, 1306–1314 (2013).

26. P Vertechi, et al., Inference-based decisions in a hidden state foraging task: differential contributions of prefrontal cortical areas. Neuron 106, 166–176 (2020).

27. XJ Wang, Probabilistic decision making by slow reverberation in cortical circuits. Neuron 36, 955–968 (2002).

28. KF Wong, XJ Wang, A recurrent network mechanism of time integration in perceptual decisions. J. Neurosci. 26, 1314–1328 (2006).

29. HK Inagaki, et al., A midbrain-thalamus-cortex circuit reorganizes cortical dynamics to initiate movement. Cell 185, 1065–1081 (2022).

30. S Recanatesi, U Pereira-Obilinovic, M Murakami, Z Mainen, L Mazzucato, Metastable attractors explain the variable timing of stable behavioral action sequences. Neuron 110, 139–153 (2022).

31. L Logiaco, L Abbott, S Escola, Thalamic control of cortical dynamics in a model of flexible motor sequencing. Cell reports 35, 109090 (2021).

32. JH Sul, S Jo, D Lee, MW Jung, Role of rodent secondary motor cortex in value-based action selection. Nat. neuroscience 14, 1202–1208 (2011).

33. AE Rajagopalan, R Darshan, KL Hibbard, JE Fitzgerald, GC Turner, Reward expectations direct learning and drive operant matching in drosophila. Proc. Natl. Acad. Sci. 120, e2221415120 (2023).

34. SE Cavanagh, JD Wallis, SW Kennerley, LT Hunt, Autocorrelation structure at rest predicts value correlates of single neurons during reward-guided choice. elife 5, e18937 (2016).

35. B Engelhard, et al., Specialized coding of sensory, motor and cognitive variables in vta dopamine neurons. Nature 570, 509–513 (2019).

36. Z Su, JY Cohen, Two types of locus coeruleus norepinephrine neurons drive reinforcement learning. bioRxiv pp. 2022–12 (2022).

37. DJ Ottenheimer, et al., A quantitative reward prediction error signal in the ventral pallidum. Nat. neuroscience 23, 1267–1276 (2020).

38. R Darshan, A Rivkind, Learning to represent continuous variables in heterogeneous neural networks. Cell Reports 39 (2022).

39. P Miller, XJ Wang, Stability of discrete memory states to stochastic fluctuations in neuronal systems. Chaos: An Interdiscip. J. Nonlinear Sci. 16 (2006).

40. DA Braun, C Mehring, DM Wolpert, Structure learning in action. Behav. brain research 206, 157–165 (2010).

41. X Boyen, N Friedman, D Koller, Discovering the hidden structure of complex dynamic systems. arXiv preprint arXiv:1301.6683 (2013).

42. CK Starkweather, BM Babayan, N Uchida, SJ Gershman, Dopamine reward prediction errors reflect hidden-state inference across time. Nat. neuroscience 20, 581–589 (2017).

43. CK Starkweather, SJ Gershman, N Uchida, The medial prefrontal cortex shapes dopamine reward prediction errors under state uncertainty. Neuron 98, 616–629 (2018).

44. R Bartolo, BB Averbeck, Prefrontal cortex predicts state switches during reversal learning. Neuron 106, 1044–1054 (2020).

45. TP Vogels, H Sprekeler, F Zenke, C Clopath, W Gerstner, Inhibitory plasticity balances excitation and inhibition in sensory pathways and memory networks. Science 334, 1569–1573 (2011).

46. YK Wu, C Miehl, J Gjorgjieva, Regulation of circuit organization and function through inhibitory synaptic plasticity. Trends Neurosci. (2022).

47. EL Charnov, Optimal foraging, the marginal value theorem. Theor. population biology 9, 129–136 (1976).

48. DW Stephens, JR Krebs, Foraging Theory. (Prineceton University press, New Jersey), (1986).

49. BY Hayden, JM Pearson, ML Platt, Neuronal basis of sequential foraging decisions in a patchy environment. Nat. Neurosci. 14, 933–939 (2011).

50. JP Noel, et al., Multiplexed and flexible neural coding in sensory, parietal, and frontal cortices during goal-directed virtual navigation. Res. Sq. pp. 10.21203/rs.3.rs-1025042/v1 (2022).

51. ZV Guo, et al., Procedures for behavioral experiments in head-fixed mice. PloS one 9, e88678 12 (2014).

52. S Chen, et al., Brain-wide neural activity underlying memory-guided movement. BioRxiv pp. 2023–03 (2023).

53. LF Abbott, FS Chance, Drivers and modulators from push-pull and balanced synaptic input. Prog. brain research 149, 147–155 (2005).

54. A Finkelstein, et al., Attractor dynamics gate cortical information flow during decision-making. Nat. Neurosci. 24, 843–850 (2021).

55. ZV Guo, et al., Maintenance of persistent activity in a frontal thalamocortical loop. Nature 545, 181–186 (2017).

56. F Zenke, W Gerstner, S Ganguli, The temporal paradox of hebbian learning and homeostatic plasticity. Curr. opinion neurobiology 43, 166–176 (2017).

57. S Lim, et al., Inferring learning rules from distributions of firing rates in cortical neurons. Nat. neuroscience 18, 1804–1810 (2015).

58. U Pereira, N Brunel, Attractor dynamics in networks with learning rules inferred from in vivo data. Neuron 99, 227–238 (2018).

59. KD Miller, F Fumarola, Mathematical equivalence of two common forms of firing rate models of neural networks. Neural computation 24, 25–31 (2012).

60. M Watabe-Uchida, L Zhu, SK Ogawa, A Vamanrao, N Uchida, Whole-brain mapping of direct inputs to midbrain dopamine neurons. Neuron 74, 858–873 (2012).

