## Supplemental Information for "Brain mechanism of foraging: reward-dependent synaptic plasticity or neural integration of values?"

#### This PDF file includes:

Supporting text
Figs. S1 to S4
SI References

### Supporting Information Text

#### Synaptic model.

***Fixed-point.*** The model in Eqs. (13, 14) in the Methods has a single fixed-point attractor that corresponds to the stationary state of the dynamics. For calculating its fixed-point, let us first define the following trial-dependent matrix

$$J(k) = \begin{pmatrix} w_A(k) - 1 - \frac{\rho}{2} & -\frac{\rho}{2} \\ -\frac{\rho}{2} & w_B(k) - 1 - \frac{\rho}{2} \end{pmatrix}. \qquad [1]$$

Then Eqs. (13, 14) in the Methods can be written in a vectorial form

$$\frac{d\vec{r}}{dt} = J(k)\vec{r} + \vec{I}. \qquad [2]$$

The fixed-point of Eq. (2) is given by

$$\vec{r}^*(k) = -J^{-1}(k)\vec{I}. \qquad [3]$$

The inverse is given by

$$J^{-1}(k) = \frac{1}{w_A(k)w_B(k) - (1+\frac{\rho}{2})(w_A(k) + w_B(k) + 1 + \rho} \begin{pmatrix} w_B(k) - 1 - \frac{\rho}{2} & \frac{\rho}{2} \\ \frac{\rho}{2} & w_A(k) - 1 - \frac{\rho}{2} \end{pmatrix}. \qquad [4]$$

The trial-dependent fixed-point is given by

$$r_A^*(k) = \frac{(1 + \frac{\rho}{2} - w_B(k))I_A - \frac{\rho}{2}I_B}{w_A(k)w_B(k) - (1+\frac{\rho}{2})(w_A(k) + w_B(k)) + 1 + \rho} \qquad [5]$$

$$r_B^*(k) = \frac{(1 + \frac{\rho}{2} - w_A(k))I_B - \frac{\rho}{2}I_A}{w_A(k)w_B(k) - (1+\frac{\rho}{2})(w_A(k) + w_B(k)) + 1 + \rho}. \qquad [6]$$

If $I_A = I_B = I_0$, then the fixed-points in Eqs. (5, 6) become

$$r_A^*(k) = \frac{I_0\left[1 - w_B(k)\right]}{w_A(k)w_B(k) - (1+\frac{\rho}{2})(w_A(k) + w_B(k)) + 1 + \rho} \qquad [7]$$

$$r_B^*(k) = \frac{I_0\left[1 - w_A(k)\right]}{w_A(k)w_B(k) - (1+\frac{\rho}{2})(w_A(k) + w_B(k)) + 1 + \rho}. \qquad [8]$$

The above equations show that the stationary firing rate depends on the recurrent synaptic efficacies $w_A(k)$ and $w_B(k)$. In our model, these quantities change in a trial-by-trial manner by a reward-dependent learning rule specified in the Methods.

For the calculation of the autocorrelation function, we first need to calculate the eigenvalues and eigenvectors of the matrix in Eq. (1). The eigenvalues are given by

$$\lambda_+ = \frac{1}{2}\left[w_A(k) + w_B(k) - 2 - \rho + \sqrt{\left(w_A(k) - w_B(k)\right)^2 + \rho^2}\right] \qquad [9]$$

$$\lambda_- = \frac{1}{2}\left[w_A(k) + w_B(k) - 2 - \rho - \sqrt{\left(w_A(k) - w_B(k)\right)^2 + \rho^2}\right]. \qquad [10]$$

and the eigenvectors are

$$\vec{v}_+ = \begin{bmatrix} \frac{w_B(k) - w_A(k) - \sqrt{(w_A(k) - w_B(k))^2 + \rho^2}}{\rho^2 + \left(\sqrt{(w_A(k) - w_B(k))^2 + \rho^2} + w_A(k) - w_B(k)\right)^2} \\ \frac{\rho^2}{\rho^2 + \left(\sqrt{(w_A(k) - w_B(k))^2 + \rho^2} + w_A(k) - w_B(k)\right)^2} \end{bmatrix} \qquad [11]$$

and

$$\vec{v}_- = \begin{bmatrix} \frac{w_B(k) - w_A(k) + \sqrt{(w_A(k) - w_B(k))^2 + \rho^2}}{\rho^2 + \left(\sqrt{(w_A(k) - w_B(k))^2 + \rho^2} + w_B(k) - w_A(k)\right)^2} \\ \frac{\rho^2}{\rho^2 + \left(\sqrt{(w_A(k) - w_B(k))^2 + \rho^2} + w_B(k) - w_A(k)\right)^2} \end{bmatrix}. \qquad [12]$$

Given a initial condition $\vec{r}_0$, the solution to the network dynamics in Eq. (2) with $\vec{I} = \vec{0}$ is given by

$$\vec{r}(t) = [\vec{v}_+, \vec{v}_-] \begin{pmatrix} e^{\lambda_+ t} & 0 \\ 0 & e^{\lambda_- t} \end{pmatrix} \begin{bmatrix} \vec{v}_+ \\ \vec{v}_- \end{bmatrix} \vec{r}_0. \quad [13]$$

Using the above expression and the definition of autocorrelation and autocovariance functions (1), and defining the following coefficients

$$R_+^A = \frac{\left(w_A(k) - w_B(k) + \sqrt{(w_A(k) - w_B(k))^2 + \rho^2}\right)^2}{\rho^2 + \left(\sqrt{(w_A(k) - w_B(k))^2 + \rho^2} + w_A(k) - w_B(k)\right)^2} \quad [14]$$

$$R_-^A = \frac{\left(w_B(k) - w_A(k) + \sqrt{(w_A(k) - w_B(k))^2 + \rho^2}\right)^2}{\rho^2 + \left(\sqrt{(w_A(k) - w_B(k))^2 + \rho^2} + w_B(k) - w_A(k)\right)^2} \quad [15]$$

$$R_+^B = 1 - C_+^A \quad [16]$$

$$R_-^B = 1 - C_-^A, \quad [17]$$

we find that the autocovariance functions are given by

$$C_A(t') = \left(\frac{R_+^A}{|\lambda_+|}\right) e^{\lambda_+ t'} + \left(\frac{R_-^A}{|\lambda_-|}\right) e^{\lambda_- t'} \quad [18]$$

$$C_B(t') = \left(\frac{R_+^B}{|\lambda_+|}\right) e^{\lambda_+ t'} + \left(\frac{R_-^B}{|\lambda_-|}\right) e^{\lambda_- t'}. \quad [19]$$

Lastly, the autocorrelation functions are given by

$$ACF_A(t') = \frac{C_A(t')}{C_A(0)} \quad [20]$$

$$ACF_B(t') = \frac{C_B(t')}{C_B(0)}. \quad [21]$$

Notice that the autocorrelation functions for both populations $ACF_A(t')$ and $ACF_B(t')$ implicitly depend on the synaptic efficacies $w_A(k)$ and $w_B(k)$.

***Synaptic perturbations.*** In Fig. 6C, we study the effect of synaptic perturbations on the network's action value encoding on a version of the synaptic model in Eq. (1) in which each population is comprised of multiple neurons. In this model, there are $N$ neurons for each excitatory population. The mean of the recurrent connectivity is given by

$$\bar{J} = \begin{pmatrix} w_A \vec{1}_N \vec{1}_N^T - I & 0 \\ 0 & w_B \vec{1}_N \vec{1}_N^T - I \end{pmatrix} - \frac{\rho}{2} \vec{1}_{2N} \vec{1}_{2N}^T + X_{2N}. \quad [22]$$

The matrix is the identity matrix of dimensions $N \times N$. The vector $\vec{1}_M$ is a vector of all $M$ entries equal to 1. The matrix $X_{2N}$ has dimensions $2N \times 2N$. Its entries are independent and identically distributed (i.i.d.) as Normal random variables with mean zero and variance equal to $\sigma_X^2/N$. This matrix models the heterogeneity across the network's synaptic strengths. Importantly, once the matrix $X_{2N}$ is drawn from the Gaussian distribution, it is fixed and does not change in time.

For simulating the effect of synaptic perturbations, as in (2), we add a time-dependent random perturbation to the synaptic weights given by

$$J(t) = \bar{J} + J_R(t) \quad [23]$$

$$J_R(0) = \sigma_R Z(0) \quad [24]$$

$$J_R(t) = \lambda J_R(t-1) + \sqrt{1-\lambda}\sigma_R Z(t). \quad [25]$$

At each time, all matrix $Z(t)$ entries are i.i.d. Normal random variables with mean zero and variance equal to one. The parameter $\lambda$ controls the decay of the perturbations while the parameter $\sigma_R$ their standard deviation. Therefore, the connectivity matrix at time $t$ is given by the time-varying matrix $J(t)$.

For Fig. 6C, the parameters are $w_A = 0.8$, $w_B = 0.4$, $\rho = 0.05$, $\sigma_X = 0.15$, $\lambda = 0.1$, $\sigma_R = 0.01$, and $N = 200$.

The dynamics of the $2N$ excitatory neurons is given by

$$\frac{d\vec{r}}{dt} = J(t)\vec{r} + \vec{I}. \quad [26]$$

In Fig. S4C, we extended the model in Eq. (27) for the encoding of $p$ action values. In this scenario, its connectivity is given by

$$\bar{J} = \begin{pmatrix} w_1 \vec{1}_N \vec{1}_N^T - I & 0 & \cdots & 0 \\ 0 & w_2 \vec{1}_N \vec{1}_N^T - I & \cdots & \vdots \\ \vdots & \cdots & \ddots & 0 \\ 0 & \cdots & 0 & w_p \vec{1}_N \vec{1}_N^T - I \end{pmatrix} - \frac{\rho}{2} \vec{1}_{pN} \vec{1}_{pN}^T + X_{pN}. \qquad [27]$$

**Neural integrator.**

***RPE computation.*** As reads in Eq. (37) on the main text, the RPE is given by

$$RPE(k) = R(k) - q_x(k) \qquad x = A, B. \qquad [28]$$

Here $q_x(k)$ corresponds to the quantile of the projection of the population activity on the line attractor axis corresponding to the chosen contingency (i.e., A or B). Since the projection can have arbitrary values, the rationale for using the quantile is to normalize this value to take values between 0 and 1 to be comparable to the reward (i.e., $R \in \{0, 1\}$). The quantile is calculated as

$$q_x(k) = \text{CDF}\left(\frac{m_x(k) - \bar{m}(k)}{\bar{\sigma}(k)}\right) \qquad x = A, B. \qquad [29]$$

Here the CDF is the cumulative normal distribution. The variable $m_x(k)$ is the projection on the line attractor axis corresponding to the chosen contingency. Lastly, the variables $\bar{m}(k)$ and $\bar{\sigma}(k)$ are the running average and standard deviation of the projections, respectively. These are calculated as

$$\bar{m}(k) = \bar{m}(k-1) - \frac{1}{\tau}\left(\frac{m_A(k) + m_B(k)}{2} - \bar{m}(k-1)\right) \qquad [30]$$

$$\bar{\sigma}^2(k) = \bar{\sigma}^2(k-1)\left(1 - \frac{1}{\tau}\right) + \frac{1}{\tau}\Delta\bar{\sigma}^2(k), \qquad [31]$$

with

$$\Delta\bar{\sigma}^2(k) = \frac{1}{2}\left[(m_A(k) - \bar{m}(k))(m_A(k) - \bar{m}(k-1)) + (m_B(k) - \bar{m}(k))(m_B(k) - \bar{m}(k-1))\right]. \qquad [32]$$

In our simulations, the running average $\bar{m}(k)$ and standard deviation of the projections $\bar{\sigma}(k)$ in Eqs. (30-32) are performed in a temporal scale of approximately 50 trials, i.e., $\tau = 50$.

***Three classes of external perturbations.*** In Fig. 4, we investigate the effect of perturbations by stimulating the network for a brief period of 100ms with a constant external input $\vec{I}_{\text{ext}}$. We use three classes of external stimulation patterns with the same norm $\|\vec{I}_{\text{ext}}\| = 5$, which are summarized below.

1. The on-manifold perturbation is given by

$$\vec{I}_{\text{ext}} = \alpha \vec{v}_N + \beta \vec{v}_{N-1}. \qquad [33]$$

For Fig. 4B we use $\alpha = 5$ and $\beta = 0$.

2. The random Gaussian perturbation is given by

$$\vec{I}_{\text{ext}}(t) = \frac{A_{\text{Random}}}{\sqrt{N}} \vec{x}, \qquad [34]$$

with $x_i \overset{i.i.d.}{\sim} N(0, 1)$. For Fig. 4B we use $A_{\text{Random}} = 20$ and $A_{\text{Random}} = 5$ for Fig. S3. Notice that, the average norm of this stimulus is $\langle \|\vec{I}_{\text{ext}}\| \rangle = A_{\text{Random}}$.

3. The uniform perturbation is given by

$$\vec{I}_{\text{ext}} = \frac{-A_{\text{Uniform}}}{\sqrt{N}} \vec{1}, \qquad [35]$$

where $\vec{1}$ is the vector with all its entries equal to one. We use $A_{\text{Uniform}} = 20$ for Fig. 4B and $A_{\text{Uniform}} = 5$ for Fig. S3.

***Connectivity perturbation.*** In Fig. S4A,B, we explore the effect of random connectivity perturbations on the action value maintenance of multiple action values (see Eq. (28) in the Methods). We perturb the neural integrator's connectivity $\mathbf{L}$ with a random Normal perturbation $\mathbf{X}_p$. The entries of this matrix are i.i.d. Normal with mean zero and variance equal to $\sigma_p^2/N$. The final connectivity is given by

$$\mathbf{L}_\mathrm{p} = \mathbf{L} + \mathbf{X}_p. \qquad [36]$$

The dynamics of the perturbed neural integrator are given by

$$\frac{d\vec{r}}{dt} = \mathbf{L}_\mathrm{p}\vec{r} + \vec{I}. \qquad [37]$$

For Fig. S4A, B, $\epsilon = -0.001$ and $\sigma_p = 0.003$.

***Engineered perturbations.*** For constructing the engineered perturbations in Fig. 4E-G we first ranked the entries of the line attractor vector that will be used for the external stimulation $\vec{v}_N$ as $v_{l_1}, v_{l_2}, \cdots, v_{l_N}$, where $v_{l_1}$ has the largest positive value and $v_{l_N}$ the largest negative value. Higher ranked neurons (e.g., neuron $l_1$) will strongly encode action value for the contingency encoded by $\vec{v}_N$. Then we constructed several stimuli of the form

$$\vec{I}_\mathrm{ext}(K) = \frac{A_\mathrm{Engineered}}{\sqrt{\sum_{i=1}^{K} v_{l_i}^2}}\left(0, \cdots, v_{l_1}, 0, \cdots, v_{l_2}, 0, \cdots, v_{l_K}, 0, \cdots\right)^T. \qquad [38]$$

Here the stimulation vector $\vec{I}_\mathrm{ext}(K)$ has only non-zero entries for the corresponding first $K$ ranked entries. For the network with $N = 1000$ neurons in Fig. 4E-G, we use $K = 1, 10, 100, 1000$ and $A_\mathrm{Engineered} = 4$.

#### Fitting network models using behavioral data.

***Reduced network models.*** For the fitting of the neural integrator and synaptic network models on behavioral data, as in (3), we first reduced the action selection network for selecting the choice $c \in \{A, B\}$ to Bernoulli random variable with probability $p(c)$. The function $p(c)$ corresponds to a sigmoidal function of the difference of the input currents that project to the action selection network from the valuation network, i.e., the Softmax function given by

$$p(c) = \begin{cases} \frac{1}{1+\exp(-\beta(I_A - I_B)+\alpha)} & \text{if } c = A \\ 1 - \frac{1}{1+\exp(-\beta(I_A - I_B)+\alpha)} & \text{if } c = B. \end{cases} \qquad [39]$$

Here $\alpha$ and $\beta$ are two parameters fitted from behavior.

From Eqs. (19-22) in the Methods for the synaptic model, the synaptic weights change according to the below equations.

$$w_{ch}(k+1) = \rho_w w_{ch}(k) + A_H(R(k) - [r^*_{ch}(k)]_+) \qquad ch \in \{A, B\} \qquad [40]$$

and for the unchosen site

$$w_{uch}(k+1) = \rho_w w_{uch}(k) \qquad uch \in \{A, B\}. \qquad [41]$$

The term $r^*_{ch}(k)$ is the valuation network's stationary firing rate of the chosen population given by Eqs. (7, 8). The function $[\bullet]_+$ is the rectifier linear function. Lastly, the term $[r^*_{ch}(k)]_+$ correspond to the predicted reward (4) decoded directly from the valuation network.

The input currents to the action selection softmax in Eq. (39) are given by

$$I_A = j_A r^*_A(k) \qquad [42]$$
$$I_B = j_B r^*_B(k) \qquad [43]$$

Here $j_A = j_B = 1$. We approximate the firing rates after the go cue by the stationary firing rates $r^*_A(k)$ and $r^*_B(k)$ at trial $k$ given by Eqs. (7, 8).

For the neural integrator, by solving the dynamics during the ITI in Eqs.(31, 32) in the Methods we found that at the time of go cue $T(k)$, the projection to the action value axis for the chosen contingency is given by

$$m_{ch}(k+1) = \left[m_{ch}(k) + A_\mathrm{RPE}(R(k) - q_{ch}(k))\right] e^{-\epsilon T(k)} \qquad ch \in \{A, B\}, \qquad [44]$$

and for the unchosen one

$$m_{uch}(k+1) = m_{uch}(k) e^{-\epsilon T(k)} \qquad ch \in \{A, B\}. \qquad [45]$$

Notice that $\epsilon$ corresponds to the drift parameter in Eq. (27) in the Methods. The input currents to the action selection softmax in Eq. (39) are given by

$$I_A = j_A m_A(k) \quad [46]$$

$$I_B = j_B m_B(k). \quad [47]$$

We chose $j_A = j_B = 1$.

***Rescorla-Wagner model.*** We also fitted the Rescorla-Wagner model (5) in the mouse foraging behavior. In this model, for a given trial $k$, the value of the chosen site is updated by

$$V_{ch}(k+1) = V_{ch}(k) + A_{RW}(R(k) - V_{ch}(k)), \quad [48]$$

while the value of the unchosen site is not updated.

For producing the choice, the choice probability is given by the softmax function in Eq. (39) with $I_A = V_A$ and $I_B = V_B$.

***Fitting procedure.*** Then the likelihood is given by

$$p(\{\text{choices}, \text{rewards}\}|\text{Parameters}) = \prod_{k=1}^{N_T} p(c_k|m_R(k), m_L(k), \text{Parameters}) \quad [49]$$

where $c_k$ is the animal's actual choice on trial $k$ and $N_T$ is the total trial number. We fitted the parameters for the three models by maximizing this likelihood using differential evolution (python package **scipy.optimize**). For the behavioral session shown in Fig. 1E, $N_T = 1,067$.

For Fig. 1E, the fitted parameters are specified below.

- For the neural integrator are $\beta = 95.5$ (Eq. (39)), $\alpha = -0.44$ (Eq. (39)), $A_{\text{RPE}} = 0.01$ (Eqs. (44, 45)), and $\epsilon = 0.042/s$ (Eqs. (44, 45)).
- For the synaptic model are $\beta = 100$ (Eq. (39)), $\alpha = -0.48$ (Eq. (39)), $A_H = 1$ (Eqs. (40, 41)), $\rho_w = 0.19$ (Eqs. (40, 41)), $I_0 = 0.09$ (Eqs. (7, 8)), and $\rho = 0.11$ (Eqs. (7, 8)).
- For the Rescorla–Wagner model (5), the fitted learning rate was 0.15. For producing the choice we use the Softmax in Eq. (39) with the fitted parameters $\beta = 4.16$ (Eq. (39)) and $\alpha = -0.89$ (Eq. (39)).

### Code

The code for simulating the network models is available at the GitHub repository https://github.com/ulisespereira/foraging-integrator-vs-synaptic. The code for fitting the reduced network models on mice behavior is available upon request.

### References


1. RN Bracewell, RN Bracewell, *The Fourier transform and its applications.* (McGraw-Hill New York) Vol. 31999, (1986).
2. M Gillett, U Pereira, N Brunel, Characteristics of sequential activity in networks with temporally asymmetric hebbian learning. *Proc. Natl. Acad. Sci.* **117**, 29948–29958 (2020).
3. A Soltani, XJ Wang, A biophysically based neural model of matching law behavior: melioration by stochastic synapses. *J. Neurosci.* **26**, 3731–3744 (2006).
4. RS Sutton, AG Barto, *Reinforcement learning: An introduction.* (MIT press), (2018).
5. RA Rescorla, A theory of pavlovian conditioning: Variations in the effectiveness of reinforcement and nonreinforcement. *Curr. research theory* pp. 64–99 (1972).

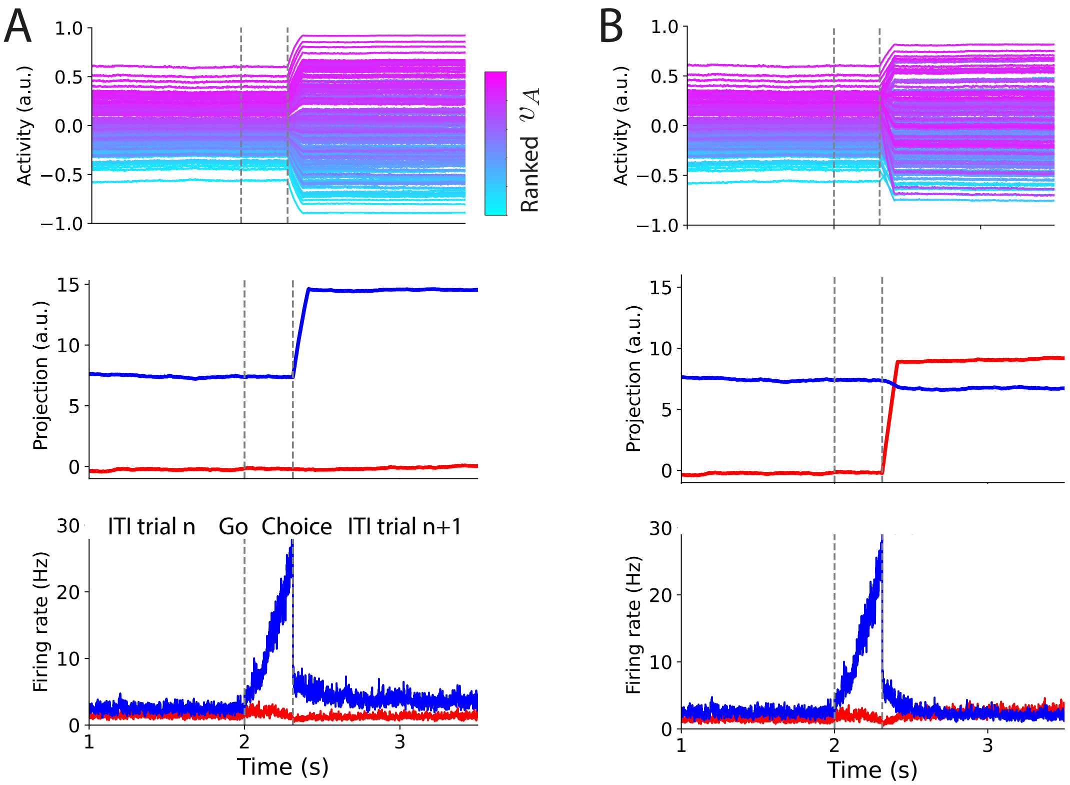


**Fig. S1.** Single trial dynamics for the neural integrator. (Top row) The activity of 1000 neurons during a single trial. Traces are color-coded according to the value of the corresponding entries of the first contingency's line attractor axis. (Middle row) The dynamics of the projections of the Valuation network's activity on the line two attractors axes. (Bottom row) The dynamics of the selective populations on the action selection network. (A) and (B) correspond to two different values of the RPE signal.

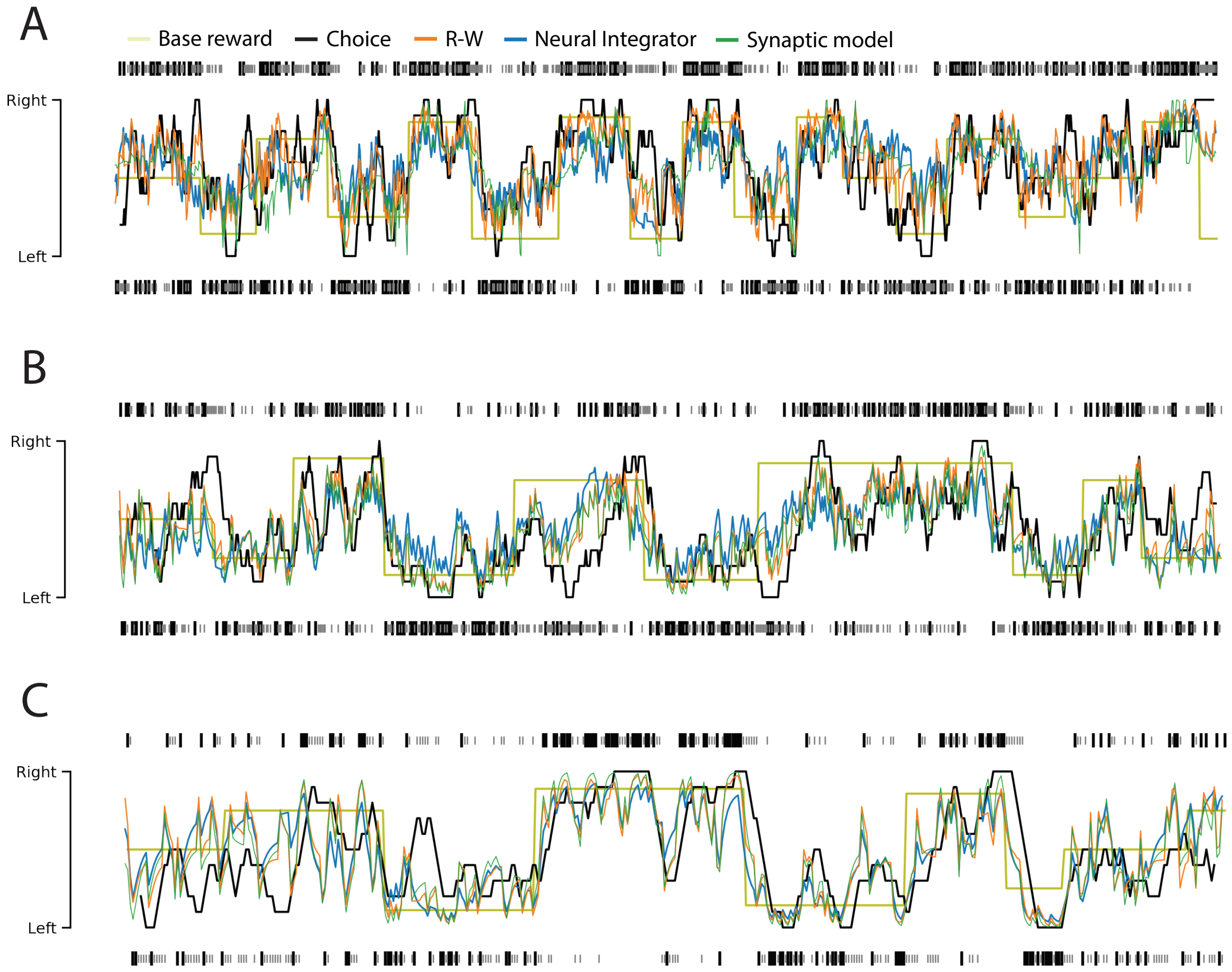


**Fig. S2.** Foraging behavior for a mouse vs. network models during an example DFT session for three different sessions (A-C). The base reward probability (pale green) changes in blocks. Black and gray ticks indicate the rewarded and unrewarded trials respectively for right (top) and left (bottom) choices. Black trace: the proportion of choices smoothed using a running average of 10 trials. The behavior was fitted using a Rescorla–Wagner model (5) (R-W) (orange), the neural integrator (blue), and a synaptic model (green). For the sessions in panels in A-C the fitted parameters are respectively: For the neural integrator $\beta = 96.4, 8.9, 29.1$ (Eq. (39)), $\alpha = -0.38, 0.31, -0.45$ (Eq. (39)), $A_{\text{RPE}} = 0.011, 0.21, 0.08$ (Eqs. (44, 45)), and $\epsilon = 0.040, 0.024, 0.055$ 1/s (Eqs. (44, 45)); For the synaptic model $\beta = 6.99, 100, 100$ (Eq. (39)), $\alpha = -0.332, 0.37, -0.50$ (Eq. (39)), $A_H = 0.99, 1, 0.582$ (Eqs. (40, 41)), $\rho_w = 0.027, 0.82, 0.20$ (Eqs. (40, 41)), $I_0 = 0.19, 0.20, 0.26$ (Eqs. (7, 8)), and $\rho = 0.11, 0.82, 1.0$ (Eqs. (7, 8)); For the Rescorla–Wagner model (5), the fitted learning rates were $0.26, 0.28, 0.126$, $\beta = 2.63, 3.57, 6.02$ (Eq. (39)) and $\alpha = -0.59, 0.46, -0.94$ (Eq. (39)).

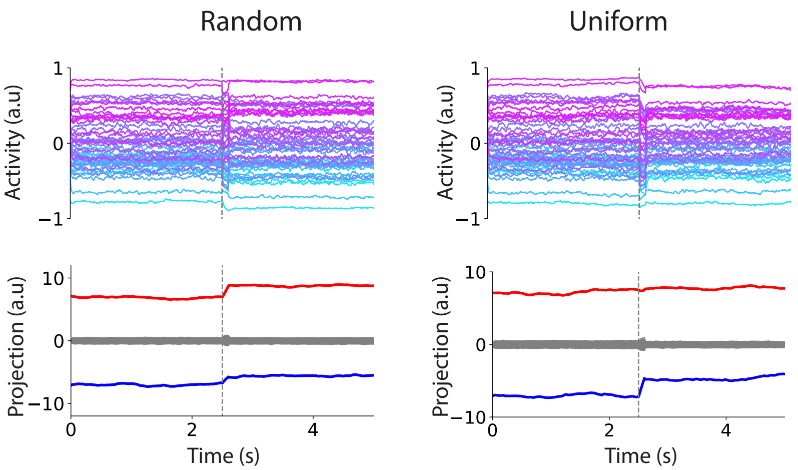


**Fig. S3.** Examples of random and uniform brief (100ms) perturbations (at the vertical dashed lines) in the valuation network for the neural integrator. Top panel: Activity of 50 representative neurons color-coded according to the magnitude of the corresponding entries of the first line attractor axis $\vec{v}_A$. Bottom panel: Projections on the line attractor axes. The magnitude of the perturbation is the same as in the on-manifold perturbation in Fig. 5B. For the stimuli parameters see the subsection "Three classes of external perturbations" on the SI text.

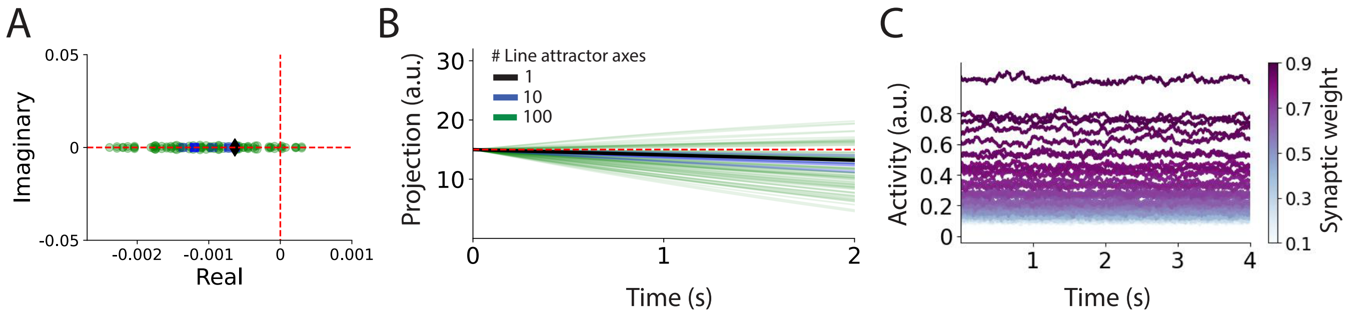


**Fig. S4.** Response to synaptic perturbations when multiple action values are encoded: Synaptic model vs. Neural integrator. (A) Zoom to the eigenvalue spectra of the perturbed neural integrator connectivity $\mathbf{L}_p$ (see Eq. (36) on the SI) with $\epsilon = -0.001$ and $\sigma_p = 0.003$. The black diamonds, blue squares, and green circles markers correspond to neural integrators with 1, 10, 1000 line attractor axes respectively. (B) Exponential drift of the network activity $\vec{u}$ projected on the line attractor axes for neural integrators with 1 (black), 10 (blue), and 100 (green) line attractor axes. (C) Average activity across $N = 50$ neurons of $p = 60$ different selective populations (see Eq. (26) in SI). The excitatory recurrent connectivity strength for each excitatory population $i$ is linearly arranged to be between 0.1 (light blue) and 0.9 (dark purple), i.e., $w_i = 0.1 + 0.8(i-1)/p \quad i = 1, \cdots, p$. The rest of the parameters are the same as those used in Fig. 6 in the main text.
